# An atlas of gene expression variation across the *Caenorhabditis elegans* species

**DOI:** 10.1101/2022.02.06.479320

**Authors:** Gaotian Zhang, Nicole M. Roberto, Daehan Lee, Steffen R. Hahnel, Erik C. Andersen

## Abstract

Phenotypic variation in diverse organism-level traits have been studied in *Caenorhabditis elegans* wild strains, but differences in gene expression and the underlying variation in regulatory mechanisms are largely unknown. Here, we use natural variation in gene expression to connect genetic variants to differences in organismal-level traits, including drug and toxicant responses. We performed transcriptomic analysis on 207 genetically distinct *C. elegans* wild strains to study natural regulatory variation of gene expression. Using this massive dataset, we performed genome-wide association mappings to investigate the genetic basis underlying gene expression variation and revealed complex genetic architectures. We found a large collection of hotspots enriched for expression quantitative trait loci across the genome. We further used mediation analysis to understand how gene expression variation could underlie organism-level phenotypic variation for a variety of complex traits. These results reveal the natural diversity in gene expression and possible regulatory mechanisms in this keystone model organism, highlighting the promise of gene expression variation in shaping phenotypic diversity.

## Introduction

Quantitative genetic mapping approaches, such as genome-wide association (GWA) and linkage mapping, have been used in a variety of organisms to disentangle the underlying genetic basis of gene expression variation by considering the expression level of each gene as a quantitative trait^1–9^. Expression quantitative trait loci (eQTL) affecting gene expression are often classified into local eQTL (located close to the genes that they influence) and distant eQTL (located further away from the genes that they influence)^10, 11^. Local eQTL are abundant in the genome. For example, over half the genes in yeast and 94.7% of all protein-coding genes in human tissues are hypothesized to have associated local eQTL^7, 8^. Genetic variants underlying local eQTL might influence the expression of a specific gene by affecting transcription factor binding sites, chromatin accessibility, other promoter elements, enhancers, or other factors at post-transcriptional levels^12^. Genes encoding diffusible factors, such as transcription factors, chromatin cofactors, and RNAs, are often considered the most likely genes to underlie distant eQTL. Distant eQTL hotspots in several species have been suggested to account for the variation in expression of many genes located throughout the genome^2, 3, 7, 9, 13^. Although a substantial amount of eQTL have been identified in different species, it is still largely unknown how gene expression variation relates to organism-level phenotypic differences.

The nematode *Caenorhabditis elegans* is a powerful model to study the genetic basis of natural variation in diverse quantitative traits^14–16^. Genome-wide gene expression variation in different developmental stages and various conditions at the whole-organism or cellular resolution have been discovered and thousands of eQTL have been identified in several studies over the past two decades^3, 9, 17–23^. However, most of these studies used two-parent recombinant inbred lines derived from crosses of the laboratory-adapted reference strain, N2, and the genetically diverse Hawaiian strain, CB4856. Consequently, the observed variation in gene expression and their identified eQTL were limited to the differences among a small number of *C. elegans* strains and only revealed a tiny fraction of the natural diversity of gene expression and regulatory mechanisms in this species. The *C. elegans* Natural Diversity Resource (CeNDR) has a collection of 540 genetically distinct wild *C. elegans* strains^16, 24, 25^. Variation in diverse organism-level phenotypes has been observed among these wild strains, and many underlying QTL, quantitative trait genes (QTGs), and quantitative trait variants (QTVs) have been identified using GWA mappings^15, 16^. Therefore, a genome-wide analysis could improve our understanding of the role of gene regulation in shaping organism-level phenotypic diversity, adaptation, and evolution of *C. elegans*.

Here, we investigated the natural variation in gene expression of 207 genetically distinct *C. elegans* wild strains by performing bulk mRNA sequencing on synchronized young adult hermaphrodites. We used GWA mappings to identify 6,545 eQTL associated with variation in expression of 5,291 transcripts of 4,520 genes. We found that local eQTL explained most of the narrow-sense heritability and showed larger effects on expression variation than distant eQTL. We identified 67 hotspots that comprise 1,828 distant eQTL across the *C. elegans* genome. We further found a diverse collection of potential regulatory mechanisms that underlie these distant eQTL hotspots. Additionally, we applied mediation analysis to gene expression and other quantitative trait variation data to elucidate putative mechanisms that can play a role in organism-level trait variation. Our results provide an unprecedented resource of transcriptome profiles and genome-wide regulatory regions that facilitate future studies. Furthermore, we demonstrate efficient methods to locate causal genes that underlie mechanisms of organism-level trait differences across the *C. elegans* species.

## Results

### Transcriptome profiles of 207 wild *C. elegans* strains

We obtained 207 wild *C. elegans* strains from CeNDR^25^ (Fig. 1a). We grew and harvested synchronized populations of each strain at the young adult stage in independently grown and prepared biological replicates (Fig. 1b). We performed bulk RNA sequencing to measure expression levels and aligned reads to strain-specific transcriptomes (Fig. 1b, Supplementary Fig. 1, Supplementary Data 1). We focused on protein-coding genes and pseudogenes and filtered out those genes with low and/or rarely detected expression (See Methods). Because various hyper-divergent regions with extremely high nucleotide diversity were identified in the genomes of wild *C. elegans* strains^26, 27^, RNA sequencing reads might be poorly aligned and expression abundances might be underestimated for genes in these regions. For each strain, we filtered out transcripts that fell into the known hyper-divergent regions. We also dropped outlier samples by comparing sample-to-sample expression distances (Supplementary Fig. 1). To further verify the homogeneity of developmental stages of our samples, we evaluated the age of each sample when they were harvested using our expression data and published time-series expression data^28^. We inferred that our animals fit an expected developmental age of 60 to 72 hours post hatching (Fig. 1c), during which time the animal is in the young adult stage. Because we harvested the animals at the first embryo-laying event, the age estimation also reflects natural variation in the duration from hatching to the beginning of embryo-laying of wild *C. elegans*. In summary, we obtained reliable expression abundance measurements for 25,849 transcripts from 16,094 genes (15,364 protein-coding genes and 730 pseudogenes) in 561 samples of 207 *C. elegans* strains (Fig. 1b, Supplementary Fig. 1, Supplementary Data 1), which we used for downstream analyses.

**Fig. 1:**
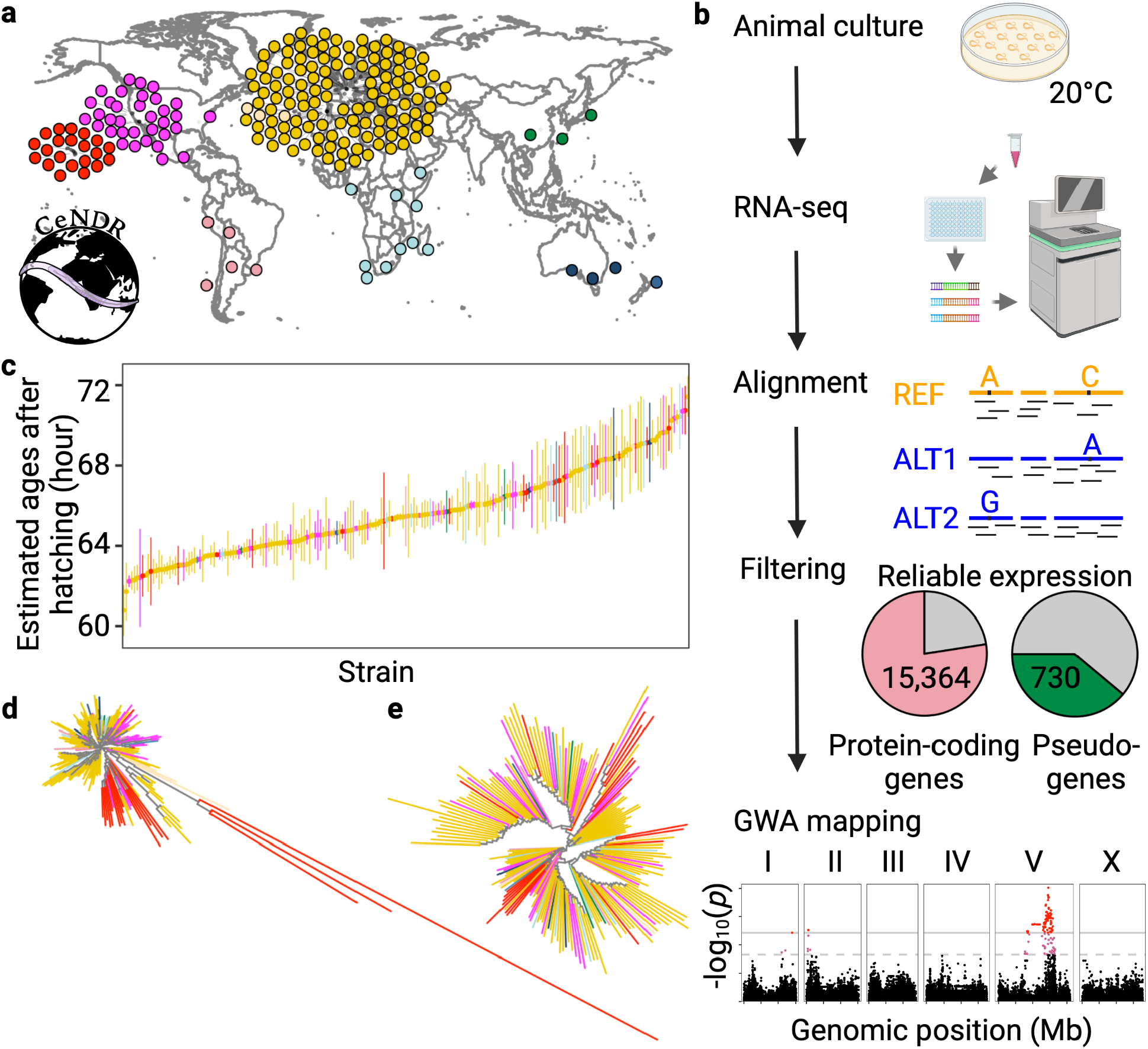
Overview of species-wide expression analysis in wild *C. elegans*. **a** Global distribution of 205 of the 207 wild *C. elegans* strains that were obtained from CeNDR and used in this study. Strains are colored by their sampling location continent, except for Hawaiian strains (in red). The two strains missing on the map are lacking sampling locations. **b** Graphic illustration of the workflow to acquire *C. elegans* transcriptome data. Created using BioRender.com. **c** Estimated developmental age (y-axis) of 561 well clustered samples of the 207 wild *C. elegans* strains (x-axis). Strains on the x-axis are sorted by their mean estimated age from two to three biological replicates. Error bars show standard deviation of estimated age among replicates of each strain. **d, e** Two Neighbor-joining trees of the 207 *C. elegans* strains using 851,105 biallelic segregating sites (**d**) and expression of 22,268 transcripts (**e**) are shown. Strains in **c, d, e** are colored as in **a**.

*C. elegans* geographic population structure has been observed previously^24, 27, 29^. Wild strains from Hawaii and other regions in the Pacific Rim harbor high genetic diversity and group into distinct clusters using genetic relatedness and principal component analysis^24, 27, 29^. Other strains that were isolated largely from Europe have relatively low genetic diversity because of the recent selective sweeps^24, 27, 29^. Similar to the previous results, the 207 strains were classified into three major groups consisting of strains from Hawaii, the Pacific coast of the United States, and Europe, respectively, in the genetic relatedness tree (Fig. 1d). Three Hawaiian strains are extremely divergent from all other strains. However, a tree constructed using transcriptome data only exhibited weak geographic relationships and no highly divergent strains (Fig. 1e), suggesting stabilizing selection has constrained variation in gene expression.

### Complex regulatory genetic architectures in wild *C. elegans* strains

To estimate the association between gene expression differences and genetic variation, we calculated the broad-sense heritability (*H*^2^) and the narrow-sense heritability (*h*^2^) for each of the 25,849 transcript expression traits. We observed a median *H*^2^ of 0.31 and a median *h^2^* of 0.06 (Fig. 2a, Supplementary Data 1), indicating strong influences from environmental factors, epistasis, or other stochastic factors on transcript expression variation^7, 30, 31^. Nearly 4,000 traits have a *h*^2^ higher than 0.18, indicating a substantial heritable genetic component of the population-wide expression differences. We performed marker-based GWA mappings to investigate the genetic basis of expression variation in the 25,849 transcripts (Supplementary Data 1). We determined the 5% false discovery rate (FDR) significance threshold for eQTL detection by mapping 40,000 permuted transcript expression traits using the EMMA algorithm^32^ and the eigen- decomposition significance (EIGEN) threshold^33^ (See Methods). In total, we detected 6,545 significant eQTL associated with variation in expression of 5,291 transcripts from 4,520 genes (Fig. 2b, Supplementary Data 2). The correlation of *h^2^* and *H^2^* among traits with eQTL is much higher than among traits without eQTL (Kendall’s τ coefficient, 0.45 and 0.27, respectively) (Fig. 2a), indicating major roles of additive genetic variation on expression variation than other genetic factors. Likely because GWA mappings mainly detect QTL that contribute additively to trait variance, eQTL were detected for 71% of the traits with *h^2^* > 0.18, but only 11% of the remaining traits (Fig. 2a).

**Fig. 2:**
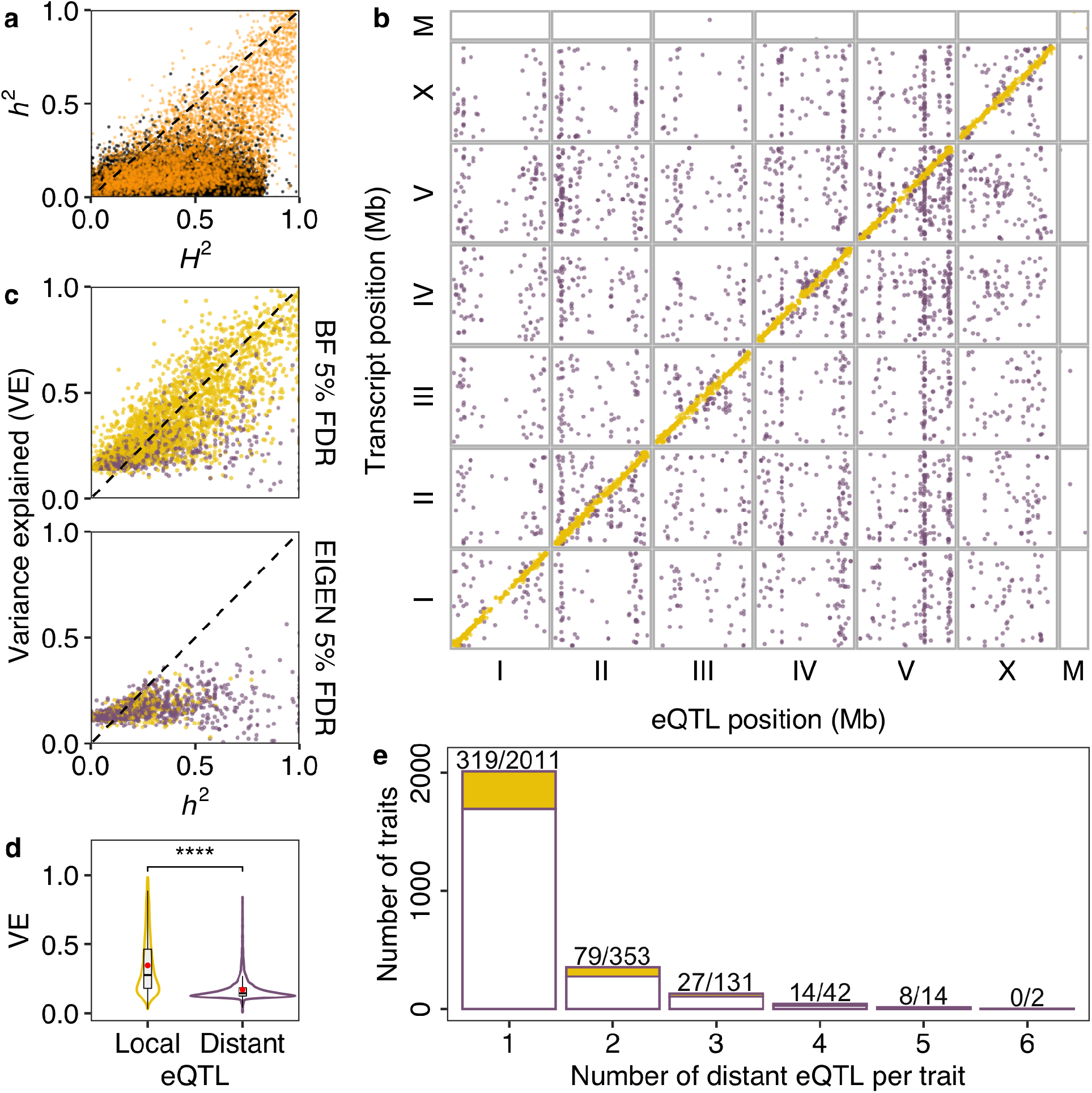
Expression QTL map of 207 wild *C. elegans* strains. **a** Heritability for 25,849 transcript expression traits with (orange) or without (black) detected eQTL. The narrow-sense heritability (*h^2^*, y-axis) for each trait is plotted against the broad-sense heritability (*H^2^*, x-axis). **b** The genomic locations of 6,545 eQTL peaks (x-axis) that pass the genome-wide EIGEN 5% FDR threshold are plotted against the genomic locations of the 5,291 transcripts with expression differences (y-axis). Golden points on the diagonal of the map represent local eQTL that colocalize with the transcripts that they influence. Purple points correspond to distant eQTL that are located further away from the transcripts that they influence. **c** The variance explained (VE) by each detected eQTL (y-axis) that passed Bonferroni (BF) 5% FDR or EIGEN 5% FDR threshold for each trait is plotted against the narrow-sense heritability *h^2^* (x-axis). The dashed lines on the diagonal are shown as visual guides to represent *h^2^* = *H^2^* (**a**) and VE = *h^2^* (**c**). **d** Comparison of VE between detected local and distant eQTL shown as Violin plots. The mean VE by local or distant eQTL is indicated as red points. Statistical significance was calculated using the Wilcoxon test with *p-value* < 2e-16 indicated as ****. **e** A histogram showing the number of distant eQTL detected per transcript expression trait. One to six distant eQTL were detected for 2,553 transcript expression traits, of which 447 traits also have one local eQTL. Numbers before slashes (indicated as the golden proportion of each bar) represent the number of traits with a local eQTL in addition to their distant eQTL. Numbers after each slash on top of each bar represent the total number of traits in each category.

In close agreement to previous *C. elegans* eQTL studies using recombinant inbred advanced intercross lines (RIAILs) derived from a cross of the N2 and CB4856 strains^3, 9^, eQTL in this study were mostly found on chromosome arms (61%) relative to centers (33%), which is likely related to the genomic distribution of genomic variation (Table 1). Of the 4,520 genes with transcript-level eQTL, we found overrepresentation of nonessential genes (Fisher exact test, odds ratio: 1.18, *p*-value: 0.001) and underrepresentation of essential genes (Fisher exact test, odds ratio: 0.75, *p*-value: 0.001), suggesting stronger selection against expression variation in essential genes than nonessential genes^34^. Gene set enrichment analysis (GSEA) on these 4,520 genes showed that proteolysis proteasome-related genes (Fisher Exact Test, Bonferroni FDR corrected *p* = 3.76E-20), especially genes encoding E3 ligases containing an F-box domain (Fisher Exact Test, Bonferroni FDR corrected *p* = 3.73E-15), are the most significantly enriched class (Supplementary Fig. 2, Supplementary Data 3). Other significantly enriched gene classes include metabolism (Fisher Exact Test, Bonferroni FDR corrected *p* = 2.92E-12), stress response (Fisher Exact Test, Bonferroni FDR corrected *p* = 7.24E-12), and histones (Fisher Exact Test, Bonferroni FDR corrected *p* = 3.23E-8). (Supplementary Fig. 2).

**Table 1:**
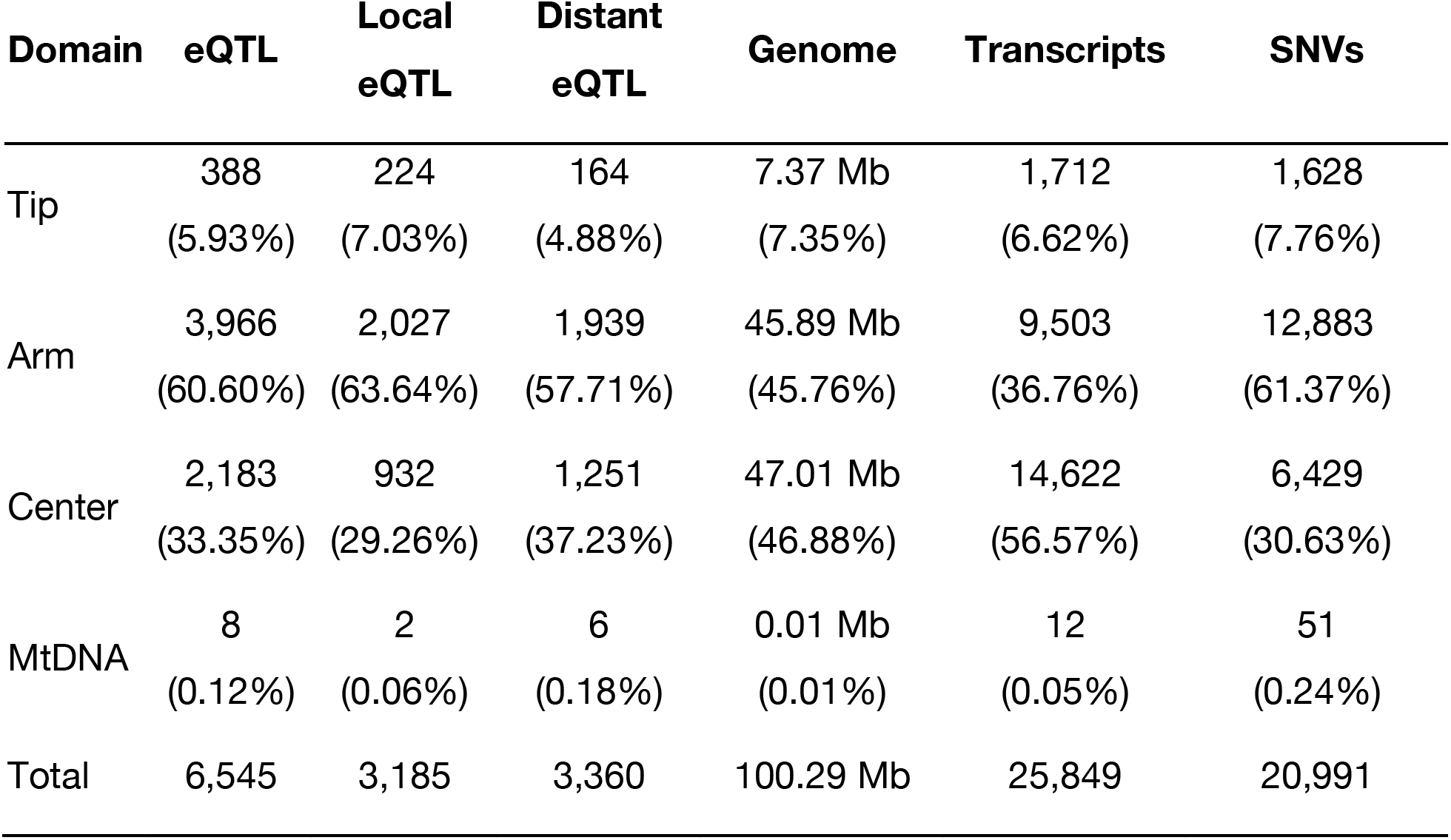
The distribution of eQTL and SNVs. Genomic domain coordinates were defined previously^36^. Transcript expression traits and SNVs used for eQTL mappings were listed.

| Domain | eQTL | Local eQTL | Distant eQTL | Genome | Transcripts | SNVs |
| --- | --- | --- | --- | --- | --- | --- |
| Tip | 388<br>(5.93%) | 224<br>(7.03%) | 164<br>(4.88%) | 7.37 Mb<br>(7.35%) | 1,712<br>(6.62%) | 1,628<br>(7.76%) |
| Arm | 3,966<br>(60.60%) | 2,027<br>(63.64%) | 1,939<br>(57.71%) | 45.89 Mb<br>(45.76%) | 9,503<br>(36.76%) | 12,883<br>(61.37%) |
| Center | 2,183<br>(33.35%) | 932<br>(29.26%) | 1,251<br>(37.23%) | 47.01 Mb<br>(46.88%) | 14,622<br>(56.57%) | 6,429<br>(30.63%) |
| MtDNA | 8<br>(0.12%) | 2<br>(0.06%) | 6<br>(0.18%) | 0.01 Mb<br>(0.01%) | 12<br>(0.05%) | 51<br>(0.24%) |
| Total | 6,545 | 3,185 | 3,360 | 100.29 Mb | 25,849 | 20,991 |

We classified eQTL located within a two megabase region surrounding each transcript as local eQTL and all other eQTL as distant^3, 9^ (Fig. 2b, Table 1, Supplementary Data 2). We identified local eQTL for 3,185 transcripts from 2,655 genes (Fig. 2b, Table 1, Supplementary Data 2). The 2,551 local eQTL that passed the Bonferroni 5% FDR threshold explained most of the estimated narrow-sense heritability (Fig. 2c). Additionally, we found 3,360 distant eQTL for 2,553 transcripts from 2,382 genes (Fig. 2b, Table 1, Supplementary Data 2). Compared to local eQTL, distant eQTL generally explained significantly lower variance (Fig. 2c, d). We found that local eQTL and up to six distant eQTL could jointly regulate the expression of transcripts (Fig. 2e). Because substantial linkage disequilibrium (LD) is observed within ( *r^2^* > 0.6) and between ( *r^2^* > 0.2) chromosomes in wild *C. elegans* strains^24, 27, 35^, we calculated LD among eQTL of each of the 861 transcripts with multiple eQTL. We found low LD among most eQTL, with a median LD of *r^2^* = 0.19 (Supplementary Fig. 3), suggesting complex genetic architectures underlying variation in expression of these transcripts are driven by independent loci.

### Diverse nature of distant eQTL hotspots

Distant eQTL were not uniformly distributed across the genome. Of the 3,360 distant eQTL, 1,828 were clustered into 67 hotspots, each of which affected the expression of 12 to 184 transcripts (Fig. 3). Signatures of selection (Tajima’s *D* values) in hotspots are mostly negative, likely because of the recent selective sweeps (Supplementary Fig. 4)^24^.

**Fig. 3:**
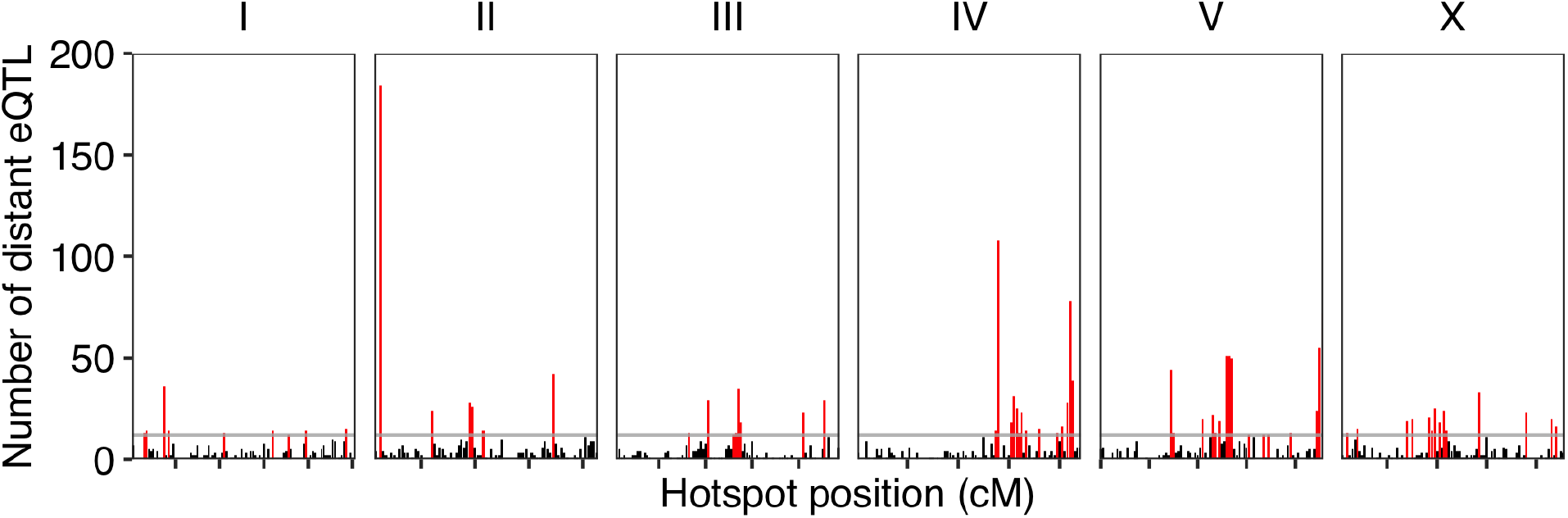
Distant eQTL hotspots. The number of distant eQTL (y-axis) in each 0.5 cM bin across the genome (x-axis) is shown. Tick marks on the x-axis denote every 10 cM. The horizontal gray line indicates the threshold of 12 eQTL. Bins with 12 or more eQTL were identified as hotspots and are colored red. Bins with fewer than 12 eQTL are colored black.

GSEA on genes with transcript-level distant eQTL in each hotspot revealed potential shared transcriptional regulatory mechanisms across different genes of the same class (Supplementary Fig. 5, Supplementary Data 3). For example, the hotspot at 21.5 cM on chromosome II significantly affected the expression of heat stress related genes (Fisher Exact Test, Bonferroni FDR corrected *p* = 7.03E-7). Our results also showed that a single hotspot could regulate expression of genes in different classes. The hotspot at 2.5 cM on chromosome II significantly affected the expression of genes in three classes, including metallopeptidases (Fisher Exact Test, Bonferroni FDR corrected *p* = 1.31E-5), collagen proteins (Fisher Exact Test, Bonferroni FDR corrected *p* = 3.11E-9), and histones (Fisher Exact Test, Bonferroni FDR corrected *p* = 1.26E-6) (Supplementary Fig. 5, Supplementary Data 3). Furthermore, different hotspots could affect the expression of the same gene class. For example, the hotspot at 45.5 cM on chromosome III was also enriched with distant eQTL of histones (Fisher Exact Test, Bonferroni FDR corrected *p* = 8.2E-7) like the hotspot at 2.5 on chromosome II (Supplementary Fig. 5, Supplementary Data 3). Regulatory genes, such as transcription factors and chromatin cofactors, that are located in each hotspot could underlie the regulation of multiple genes. We found previously known or predicted genes encoding chromatin cofactors and transcription factors^37–39^ in 24 and 59 of the 67 hotspots, respectively (Supplementary Fig. 6).

To identify causal genes and variants underlying hotspots, we performed fine mapping on distant eQTL in each hotspot and filtered for the most likely candidate variants (see Methods for details) (Supplementary Data 4). Then, we focused on the filtered candidate variants that were mapped for at least four traits in each hotspot and are in genes encoding transcription factors or chromatin cofactors. In total, we identified 36 candidate genes encoding transcription factors or chromatin cofactors for 34 hotspots. For example, the gene *ttx-1*, which encodes a transcription factor necessary for thermosensation in the AFD neurons^40, 41^, might underlie the expression variation of 97 transcripts with distant eQTL in three hotspots between 44.5 cM and 45.5 cM on chromosome V. TTX-1 regulates expression of *gcy-8* and *gcy-18* in AFD neurons^40, 41^, but no eQTL were detected for the two genes likely because we measured the expression of whole animals. Additionally, the linker histone gene *hil-2*^39^ might underlie the expression variation of 46, 10, 17, and four transcripts with distant eQTL in the hotspots at 28 cM, 30.5 cM, 31 cM and 31.5 cM, respectively, on chromosome IV. We also performed GSEA for groups of transcripts whose expression traits were fine mapped to the 36 candidate genes encoding transcription factors or chromatin cofactors. For instance, the 17 traits that fine mapped to *hil-2* in the hotspot at 31 cM on chromosome IV (Supplementary Fig. 7) were enriched in E3 ligases containing an F-box domain (Fisher Exact Test, Bonferroni FDR corrected *p* = 0.0003) and transcription factors of the homeodomain class (Fisher Exact Test, Bonferroni FDR corrected *p* = 0.002). Besides the 36 candidate genes, the hundreds of other fine mapping candidates are not as transcription factors or chromatin cofactors, suggesting other mechanisms underlying distant eQTL. Altogether, as previously implicated in other species^7, 11, 42^, our results indicate that a diverse collection of molecular mechanisms likely cause gene expression variation in *C. elegans*.

### Mediation analysis facilitates candidate gene prioritization

Mediation analysis seeks to identify the mechanism that underlies the relationship between an independent variable and a dependent variable via the inclusion of an intermediary mediating variable. Because gene expression has been found to play an intermediate role between genotypes and phenotypes, it could help to identify the causal mediating genes between genotypes and phenotypes in quantitative genetics mapping studies. We have previously identified mediation effects of *scb-1* expression on responses to several chemotherapeutics and *sqst-5* expression on differential responses to exogenous zinc using linkage mapping experiments^9, 43^. To validate if our expression and eQTL data can be used to identify candidate genes, we first performed mediation analysis on one published GWA study of variation in responses to the commonly used anthelmintic albendazole (ABZ)^44^.

Previously, wild *C. elegans* strains were exposed to ABZ and measured for effects on development to identify genomic regions that contribute to variation in ABZ resistance. A single-marker GWA mapping was performed first to detect two QTL on chromosomes II and V, but no putative candidate gene was identified. Using a burden mapping approach, prior knowledge of ABZ resistance in parasitic nematodes, and manually curation of raw sequence read alignment files, the gene *ben-1* was found to underlie natural variation in ABZ resistance variation^44^. The single-marker GWA mapping was not able to detect an association between ABZ resistance and *ben-1* variation because of high allelic heterogeneity caused by rare SNVs and structural variants (Supplementary Fig. 8). However, rare SNVs or structural variants might lead to changes in *ben-1* expression and ABZ resistance. We found two distant eQTL, in regions overlapping with the two organism-level ABZ QTL, for *ben-1* expression variation. Therefore, these results provided an excellent opportunity to test the effectiveness of mediation analysis among organism-level phenotypes, genotype, and gene expression. The mediation estimate for *ben-1* expression was the second strongest hit in the analysis on the phenotype (animal length in response to ABZ), the genotype (GWA QTL of the phenotype), and the expression of 1,157 transcripts (Fig. 4a). We found a moderate negative correlation between the expression of *ben-1* and animal length and almost no correlation after we regressed animal length by the expression of *ben-1* (Fig. 4b), suggesting that expression variation impacts differences in ABZ responses. We further examined genetic variants across strains and found that those strains with relatively low *ben-1* expression and high ABZ resistance all harbor SNVs or structural variants with different predicted effects (Fig. 4b), suggesting that the extreme allelic heterogeneity at the *ben-1* locus might affect ABZ response variation by reducing the abundance of this beta-tubulin. To test the impact of expression variation on phenotypic variation, we regressed animal length by expression of every transcript in our data and performed GWA mappings. Then, we compared the GWA mapping significance value after regression to the original GWA mapping significance value at a pseudo variant marker that represents all the variants in *ben-1* (Fig. 4c, Supplementary Fig. 8)^45^. We found animal length regressed by the expression of *ben-1* showed one of the largest drops in significance, and significance in most of the other mappings was approximately equal to the original significance value (Fig. 4c, Supplementary Fig. 8). These results indicated that increasing *ben-1* expression decreases resistance to ABZ and suggested the applicability of mediation analysis using the expression and eQTL data for other *C. elegans* quantitative traits.

**Fig. 4:**
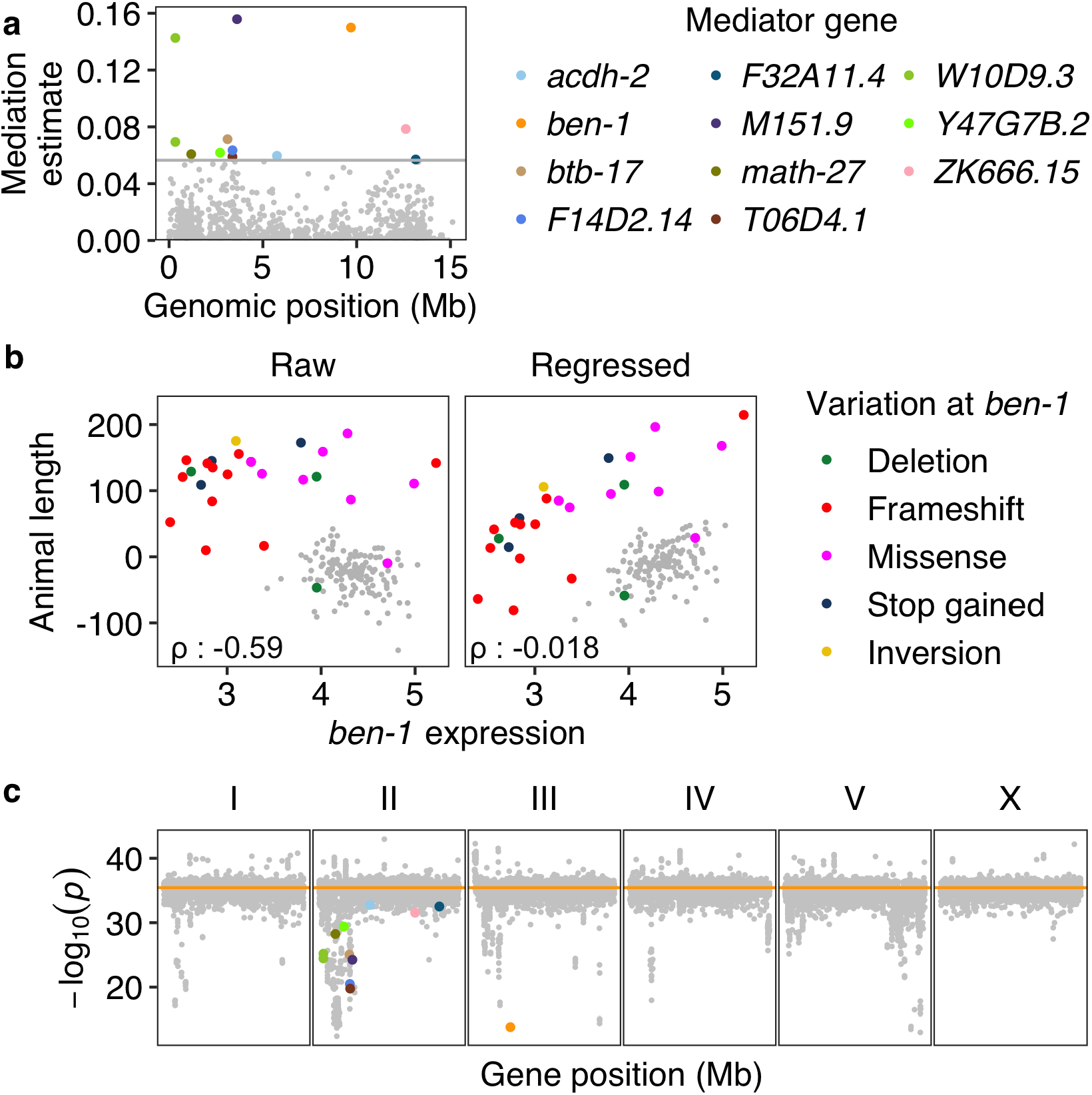
Mediation effects of *ben-1* expression on *C. elegans* resistance to ABZ. **a** Mediation estimates (y-axis) calculated as the indirect effect that differences in expression of each gene play in the overall phenotype are plotted against the genomic position of the eQTL (x-axis) on chromosome II. The horizontal gray line represents the 99^th^ percentile of the distribution of mediation estimates. Significant mediators are colored other than gray by their genes as shown in the legend. **b** The correlation of *ben- 1* expression (x-axis) to raw animal length and to animal length regressed by *ben-1* expression on y-axis. The Pearson’s coefficient ρ for each correlation was indicated at bottom left. Strains are colored by the type of their genetic variants in *ben-1*. Strains without identified variants are colored gray. **c** Significance at the pseudo variant marker of 25,837 GWA mappings. Each point represents a GWA mapping that is plotted with its -log_10_(*p*) value (y-axis) at the pseudo variant marker (III: 3,539,640) against the genomic locations (x-axis) of the transcript of which the expression was used in regression for animal length. Points for traits regressed by expression of transcripts identified as significant mediators are colored as in (**a**). The orange horizontal line represents the significance at the pseudo variant marker using the raw animal length of 167 strains (Supplementary Fig. 8). GWA mapping results of 12 traits regressed by expression of mitochondrial genes were excluded but all with significance close to the horizontal line.

We further applied mediation analysis to another eight previously published studies of *C. elegans* natural variation and GWA mappings in diverse traits, including telomere length^46^ (Fig. 5a), responses to arsenic^47^ (Fig. 5b), zinc^43^ (Fig. 5c), etoposide^48^ (Fig. 5d), propionate^49^ (Fig. 5e), abamectin^50^ (Fig. 5f), dauer formation in response to pheromone^51^, and lifetime fecundity^52^ (Fig. 5g). Causal variants and genes that partially explained the phenotypic variation in all the eight traits, except for lifetime fecundity, have been identified using fine mappings and genome-editing experiments^43, 46–52^. Only one causal gene, *dbt-1* (for arsenic response variation^47^), has eQTL detected and its expression was tested in mediation analysis for arsenic response variation^47^ (Fig. 5b). No significant mediation effects were found on arsenic response variation by the expression of *dbt-1*. We also did not observe significant differential expression between strains with different alleles at the previously validated causal *dbt-1* QTV (II:7944817)^47^. Therefore, this causal variant possibly causes arsenic response variation only by affecting enzymatic activity^47^ and not the abundance of the *dbt-1* transcript. Instead, we identified *bath-15* as a significant mediator gene for arsenic response variation (Fig. 5b). For the other seven organism-level traits, putative genes whose expression likely mediated the phenotypic variation were detected for six of the traits (Fig. 5). For example, the top mediator gene for the variation in responses to abamectin was *cyn-7*, which is predicted to have peptidyl-prolyl cis-trans isomerase acitivity (Fig. 5f)^53^. For the variation in lifetime fecundity (Fig. 5g), one of the putative mediator genes was *ets-4*, which is known to affect larval developmental rate, egg-laying rate, and lifespan^54^. Mediator genes suggest candidate genes in addition to those genes identified in fine mappings or linkage mappings. Taken together, we concluded that mediation analysis using the newly generated expression and eQTL data facilitates candidate gene prioritization in GWA studies.

**Fig. 5:**
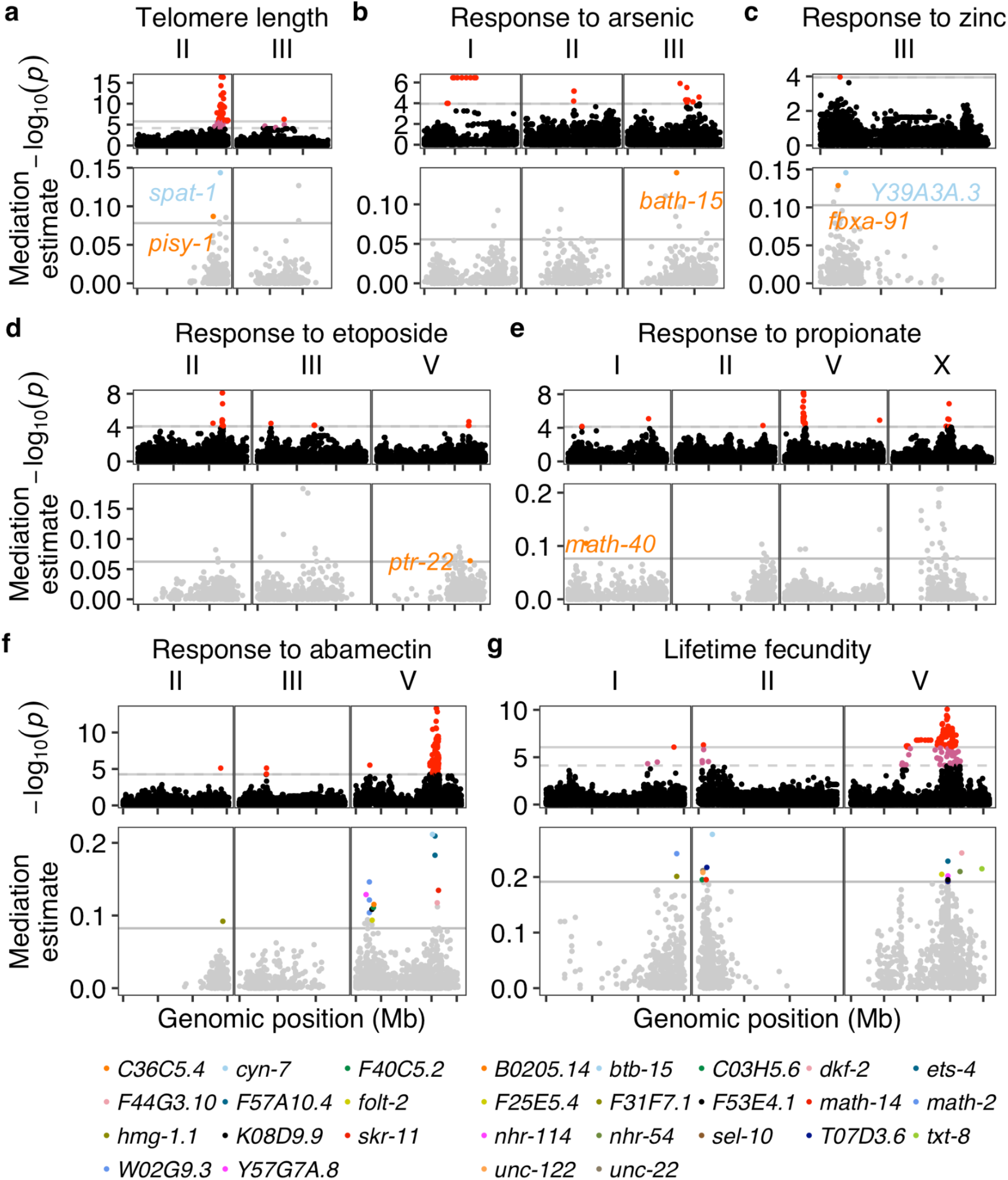
Mediation effects of gene expression on variation in seven organism-level phenotypes of *C. elegans*. GWA mapping and mediation analysis results of natural variation in *C. elegans* telomere length (**a**), responses to arsenic (**b**), zinc (**c**), etoposide (**d**), propionate (**e**), abamectin (**f**), and lifetime fecundity (**g**). Top panel: A Manhattan plot indicating the GWA mapping result for each phenotype is shown. Each point represents an SNV that is plotted with its genomic position (x-axis) against its -log_10_(*p*) value (y-axis) in mapping. SNVs that pass the genome-wide EIGEN threshold (the dotted gray horizontal line) and the genome-wide Bonferroni threshold (the solid gray horizontal line) are colored pink and red, respectively. QTL were identified by the EIGEN (**c,d,e,f**) or Bonferroni (**a,b,g**) threshold. Only chromosomes with identified QTL were shown. Bottom panel: Mediation estimates (y-axis) calculated as the indirect effect that differences in expression of each gene plays in the overall phenotype are plotted against the genomic position (x-axis) of the eQTL. The horizontal gray line represents the 99^th^ percentile of the distribution of mediation estimates. The mediator genes with adjusted *p* < 0.05 and interpretable mediation estimate > the 99^th^ percentile estimates threshold are colored other than gray and labeled in the panel (**a-e**) or below the panel (**f, g**). Tick marks on x-axes denote every 5 Mb.

## Discussion

*C. elegans* was the first metazoan to have its genome sequenced and has been subjected to numerous genetic screens to identify the genes that underlie diverse traits, including programmed cell death, drug responses, development, and behaviors. Despite huge efforts by a large research community, over 60% of its genes have not been curated with functional annotations or associated with defined mutant phenotypes^55^. A likely reason is that most *C. elegans* research uses the reference strain N2 under laboratory conditions, and the functions of many genes might only be revealed in natural environments or in different genetic backgrounds^56^. In the last decade, wild *C. elegans* strains have exhibited diverse phenotypic variation in natural ecology studies^16, 25, 29, 57–59^. Here, we provide an unprecedentedly large resource of transcriptome profiles from wild *C. elegans* strains. We used these data and GWA mappings to study gene regulation variation and detected 6,545 eQTL associated with variation in expression of 5,291 transcripts of 4,520 genes. These genes are enriched in processes, including the proteasome, metabolism, stress response, etc., suggesting gene expression regulation plays an important role in adaptation of natural *C. elegans* strains to various environments^60, 61^. We identified local eQTL that explained most of the narrow-sense heritability (*h^2^*) and significantly larger variance than distant eQTL, likely because of higher possibilities of pleiotropy and thus stronger selection pressures. We also observed lower variation in gene expression than in genome sequence and underrepresentation of essential genes among all of the genes identified with eQTL, suggesting stabilizing selection against gene expression as previously observed in *C. elegans* and other species^5, 12, 62, 63^.

Although previous *C. elegans* eQTL studies using recombinant inbred lines have revealed rich information on the genetic basis of gene expression variation, mapping using 207 genetically distinct wild strains has the advantage of much greater genetic diversity. We reanalyzed results of one previous study that used linkage mapping to identify eQTL from the young adult stage of N2xCB4856 recombinant inbred lines^3, 9^. We reclassified 1,208 local eQTL and 1,179 distant eQTL for 2,054 microarray probes of 2,003 genes (Supplementary Fig. 9a). Both the eQTL GWA and linkage mappings detected overlapping local eQTL for 454 genes and distant eQTL for 19 genes, indicating that the CB4856 strain carries the common alternative alleles among wild *C. elegans* strains for these 473 loci. However, among the 6,545 eQTL that we detected, the strains N2 and CB4856 shared the same genotypes in 4,476 eQTL, which could not be discovered using N2xCB4856 recombinant inbred lines. Alternatively, RIAILs might have less linkage disequilibrium between nearby variants and thus smaller eQTL regions of interest than eQTL in wild *C. elegans* strains. The GWA eQTL in this study have a median region of interest of 2.1 Mb (ranged from 12 kb to 18 Mb), whereas the N2xCB4856 RIAILs eQTL showed a median size of 0.55 Mb (ranged from 149 bp to 6.8 Mb), which might make the identification of underlying causal variants easier. We further found 17 distant eQTL hotspots overlapped between the two studies (Supplementary Fig. 9b). However, these shared hotspots comprise different genes between the two studies, indicating that variation in regulatory factors is not common between the linkage and association mapping studies. Future research should leverage both types of mapping studies to identify common regulatory mechanisms, focusing on local eQTL.

In addition to the high linkage disequilibrium across the *C. elegans* genome, the recently discovered hyper-divergent genomic regions made this eQTL study challenging. Approximately 20% of the genomes in some wild *C. elegans* strains were found to have extremely high diversity compared to the N2 reference genome^27^. Short-sequence reads of wild *C. elegans* strains often fail to align to the N2 reference genome in these regions and showed lower coverage than in other regions^27^. Similarly, expression levels of genes in hyper-divergent regions could be underestimated because of the poor alignment of RNA-seq reads. Therefore, we only used expression of transcripts in non-divergent regions to map eQTL and flagged the loci that are in common hyper-divergent regions, where we are less confident in the genotypes of wild strains (Supplementary Data 2). Furthermore, we only used distant eQTL that are not in common hyper-divergent regions to identify hotspots. Because hyper-divergent regions were suggested to be under long-term balancing selection, our estimates of Tajima’s *D* in hotspots are probably biased towards lower values. Future efforts using long-read sequencing are necessary to study the sequence, expression, natural selection, and evolution of genes in hyper-divergent regions.

Variation in gene expression was suggested to impact organism-level phenotypic variation^7, 64–66^. Combining previous GWA studies in *C. elegans* with expression of genes with eQTL, we used mediation analysis to search for organism-level phenotypic variation that can be explained by variation in gene expression. Compared to previous studies using mediation analysis on gene expression and eQTL data from the N2xCB4856 recombinant inbred lines^9, 43^, we added a multiple testing correction procedure to our mediation analysis. We performed mediation analysis on ABZ response variation^44^. The causal gene *ben-1* underlying the trait was identified using a burden mapping approach^44^ along with prior knowledge^67, 68^ about the role of beta-tubulin in this drug response. Although two GWA QTL on chromosomes II and V were found, they were identified likely because of their interchromosomal linkage disequilibrium to variants in the *ben-1* locus^44^ (Supplementary Fig. 8). The single-marker GWA mapping could not associate ABZ response variation because of the extreme allelic heterogeneity at the *ben-1* locus. However, we used mediation analysis to identify *ben-1* without consideration of prior knowledge or burden mapping results, demonstrating the power of the approach (Fig. 4a). We further identified significant mediators for seven other organism-level traits (Fig. 5). The expression of these mediator genes could affect the corresponding phenotypic variation, which should be validated in the future.

Mediation analysis provides an efficient hypothesis-generating approach to be performed in parallel to fine mappings. Additionally, mediator genes could contribute to organism-level phenotypic variation in addition to causal genes identified using fine mappings. One limitation of fine mappings is that searching for causal genes and variants is restricted to the QTL region of interest. Mediation analysis can make statistical connections between the organism-level phenotypes and expression of genes far away from the QTL. As mentioned above, large GWA QTL regions of interest make it difficult to identify causal genes, which require validation using genome editing. Future *C. elegans* GWA studies should use both fine mappings and mediation analysis to prioritize candidate genes. If the candidate genes overlap between the two approaches, then validation approaches can be initiated using genome editing. In cases where the two approaches identify different candidate genes, prioritization using prior knowledge across all genes identified by both approaches can inform which genes should be tested for validation using genome editing. Previous studies using fine mappings prioritized candidate genes harboring coding variants predicted to have strong functional impacts. In mediation analysis, noncoding variants that likely affect expression of mediator genes could also be nominated as candidates. For example, upstream variants were suggested to underlie expression variation of the gene *scb-1*, which mediated differences in responses to bleomycin and three other chemotherapeutics^9, 69^. To summarize, we recommend using both fine mappings and mediation analyses to nominate candidate genes and variants.

The goal of quantitative genetics is to understand the genetic basis and mechanisms underlying phenotypic variation. Here, we showed that mediation analysis, which uses expression and eQTL data to search connections between genetic variants and complex traits, provides additional loci that might further explain phenotypic variation. The framework we developed for mediation analysis complements marker-based GWA mappings and is also applicable using various other intermediate traits, such as small RNAs, proteins, and metabolites. Any genes and variants underlying variation in these factors can be nominated as candidates for phenotypic validation. Furthermore, we could measure all of these data and complex traits from the exact same samples using *C. elegans*, which can be easily grown at large scale to have synchronized isogenic populations. Analyses using measurements of mRNAs, small RNAs, proteins, and metabolites could strengthen conclusions about causal genes and mechanisms underlying complex traits using a more holistic perspective of organismal phenotypic variation. We foresee this strategy will greatly improve the powers of quantitative genetic mappings in the future.

## Methods

### C. elegans strains

We obtained 207 wild *C. elegans* strains from *C. elegans* Natural Diversity Resource (CeNDR)^25^. Animals were cultured at 20°C on modified nematode growth medium (NGMA) containing 1% agar and 0.7% agarose to prevent burrowing and fed *Escherichia coli* (*E. coli*) strain OP50^70^. Prior to each assay, strains were grown for three generations without starvation or encountering dauer-inducing conditions^70^.

### Animal growth and harvest

We grew and harvested synchronized populations of each strain at the young adult stage with independently grown and prepared biological replicates. Specifically, L4 larval stage hermaphrodites were grown to the gravid adult stage on 6 cm plates and were bleached to obtain synchronized embryos. Approximately 1,000 embryos were grown on each 10 cm plate to the young adult stage and were harvested after the first embryo was observed. M9 solution was used to wash harvested animals twice to remove *E. coli*. Animals were then pelleted by centrifugation (2000 rpm for one minute) and Trizol reagent (Ambion) was added to maintain RNA integrity before storage at -80°C.

### RNA extraction

Frozen samples in Trizol were thawed at room temperature and 100 µL acid-washed sand (Sigma, catalog no. 274739) was added to help to disrupt animal tissues. Then chloroform, isopropanol, and ethanol were used for phase separation, precipitation, and washing steps, respectively. Total RNA pellets were resuspended in nuclease-free water. The concentration of total RNA was determined using the Qubit RNA XR Assay Kit (Invitrogen, catalog no. Q33224). RNA quality was measured using the 2100 Bioanalyzer (Agilent). RNA samples with a minimum RNA integrity number (RIN) of 7 were used to construct Illumina sequencing libraries.

### RNA library construction and sequencing

Illumina RNA-seq libraries were prepared in 96-well plates. Replicates of the same strain were prepared in different 96-well plates. For each sample, mRNA was purified and enriched from 1 µg of total RNA using the NEBNext Poly(A) mRNA Magnetic Isolation Module (New England Biolabs, catalog no. E7490L). RNA fragmentation, first and second strand cDNA synthesis, and end-repair processing were performed with the NEBNext Ultra II RNA Library Prep with Sample Purification Beads (New England Biolabs, catalog no. E7775L). The cDNA libraries were adapter-ligated using adapters and unique dual indexes in the NEBNext Multiplex Oligos for Illumina (New England Biolabs, catalog no. E6440, E6442) and amplified using 12 PCR cycles. All procedures were performed according to the manufacturer’s protocols. The concentration of each RNA-seq library was determined using Qubit dsDNA BR Assay Kit (Invitrogen, catalog no. Q32853). Approximately 96 RNA-seq libraries were pooled and quantified with the 2100 Bioanalyzer (Agilent) at Novogene, CA, USA. Each of the pools of libraries were sequenced on a single lane of an Illumina NovaSeq 6000 platform, yielding 150-bp paired-end (PE150) reads.

In total, RNA-seq data of 608 samples from 207 wild *C. elegans* strains in seven pooled libraries were obtained with an average of 32.6 million reads per sample and a minimum of 16.6 million reads. Of the 207 strains, 194 strains with three replicates and 13 strains with two replicates.

### Sequence processing and expression abundance quantification

Adapter sequences and low-quality reads in raw sequencing data were removed using *fastp* (v0.20.0)^71^. *FastQC* (v0.11.8) analysis (http://www.bioinformatics.babraham.ac.uk/projects/fastqc) was performed on trimmed FASTQ files to assess read quality (adapter content, read-length distribution, per read GC content, etc.). For RNA-seq mapping, SNV-substituted reference transcriptomes for each of the wild *C. elegans* strains were generated using *BCFtools* (v.1.9)^72^, *gffread* (v0.11.6)^73^, the N2 reference genome (WS276), a GTF file (WS276)^53^, and the hard-filtered isotype variant call format (VCF) 20200815 CeNDR release (Supplementary Fig. 1).

Transposable element (TE) consensus sequences of *C. elegans* were also extracted from Dfam (release 3.3)^74^ using scripts (https://github.com/fansalon/TEconsensus). We used *Kallisto* (v0.44.0) to (1) pseudoalign trimmed RNA-seq reads from each sample to the transcriptome index built from the strain-specific SNV-substituted reference transcriptome (65,173 transcripts) and TE consensus sequences (157 TEs) and (2) quantify expression abundance at the transcript level^75^. On average, 31.3 million reads pseudoaligned to the transcriptome index per sample with a minimum of 15.5 million reads, which were sufficient to capture the expression of more than 70% of the *C. elegans* reference genome genes. We used the 608 samples of 207 strains and 39,008 transcripts of protein-coding genes and pseudogenes in our analysis.

### Selection of reliably expressed transcripts

We first normalized the raw counts of transcript expression abundances without the default filtering of low abundance transcripts using the R package *sleuth* (v0.30.0)^76^. Then, we filtered reliably expressed transcripts (26,043) by requiring at least five normalized counts in all the replicates of at least ten strains (Supplementary Fig. 1). We also filtered out 3,775 transcripts of 2,597 genes that are in hyper-divergent genomic regions of at least one strain. We further excluded 194 transcripts in hyper-divergent regions of more than 107 of the 207 strains. In summary, we collected reliable expression abundance for 25,849 transcripts of 16,094 genes (15,364 protein-coding genes and 730 pseudogenes).

### Selection of well clustered samples

We used sample-to-sample distance to select well clustered samples (Supplementary Fig. 1). We first summarized raw counts of reliably expressed transcripts into gene-level abundances using the R package *tximport* (v1.10.1)^77^. Then, we performed variance stabilizing transformations on the gene expression profile using the *vst()* function in the R package *DESeq2* (v1.26.0), which generated log_2_ scale normalized expression data^78^. Sample-to-sample pairwise Euclidean distances among the 608 samples were calculated using the generic function *dist()* in R^79^. Our basic assumption is that intra-strain distances among replicates should be smaller than inter-strain distances. Because the majority of the 207 wild *C. elegans* strains exhibit low overall genetic diversity (Fig. 1d)^24, 35, 80^, we required that the intra-strain distances of replicates be smaller than the median of inter-strain distances of the strain to other strains. Specifically, for each strain, if all of its intra-strain distances were smaller than the median of its inter-strain distances, then all of its replicates were kept. If none of its intra-strain distances were smaller than the median of its inter-strain distances, then all samples of the strain were removed. For strains with three replicates, if one or two of its three intra-strain distances were smaller than the median of its inter-strain distances, then the two replicates with the minimum distances were kept. After removal of some outlier samples, the median of inter-strain distances would change. Therefore, we repeatedly performed the procedures of data transformation, sample-to-sample distance calculation, and filtering by comparing inter- and intra-strain distances until no more samples were removed. Eventually, 561 samples of 207 strains were selected as well clustered samples, which comprised 147 strains with three replicates and 60 strains with two replicates.

### Transcript expression abundance normalization

We used the function *norm_factors()* in the R package *sleuth* (v0.30.0)^76^ to compute the normalization factors for each sample using the raw transcripts per million reads (TPM) of 22,268 reliably expressed transcripts in non-divergent regions of the 207 strains and their well clustered samples. Then, we normalized the raw TPM of all the 25,849 reliably expressed transcripts of each sample with the normalization factors and used log_2_(normalized TPM + 0.5) for downstream analysis unless indicated otherwise.

### Sample age estimation

To further verify the homogeneous developmental stage of our samples, we evaluated the age of each sample when they were harvested using the R package *RAPToR* (v1.1.3)^28^ (Supplementary Fig. 1). As the requirement of the package, we first generated gene-level expression abundances. Raw TPM of 22,268 reliably expressed transcripts in non-divergent regions were summarized into abundances of 13,637 genes using the R package *tximport* (v1.10.1)^77^. Normalization factors for each sample using gene-level abundances were calculated as described for transcript level and were used to normalize gene level TPM. Correlation of log_2_(normalized TPM + 0.5) of our data against the reference gene expression time series (Cel_YA_2) in *RAPToR* was computed using the function *ae()* in *RAPToR* with 10,489 intersected genes and default parameters.

### Genetic and expression relatedness

Genetic variation data for 207 *C. elegans* isotypes were acquired from the hard-filtered isotype variant call format (VCF) 20200815 CeNDR release. These variants were pruned to the 851,105 biallelic single nucleotide variants (SNVs) without missing genotypes. We converted this pruned VCF file to a PHYLIP file using the *vcf2phylip.py* script^81^. Expression distance among the 207 wild strains was calculated based on the mean expression of 22,268 transcripts without missing data using the function *dist()* in R. The unrooted neighbor-joining trees for genetic and expression relatedness were made using the R packages *phangorn* (v2.5.5)^82^ , *ape* (v5.6)^83^ and *ggtree* (v1.14.6)^84^.

### eQTL mapping

#### Input phenotype and genotype data

For the 25,849 transcripts, we summarized the expression abundance of replicates to have the mean expression for each transcript of each strain as phenotypes used in GWA mapping (Supplementary Data 1). Genotype data for each of the 207 strains were acquired from the hard-filtered isotype VCF (20200815 CeNDR release).

#### Permutation-based FDR threshold

We performed GWA mapping using the pipeline *cegwas2-nf* (https://github.com/AndersenLab/cegwas2-nf). The pipeline uses the eigen- decomposition significance (EIGEN) threshold or the more stringent Bonferroni-corrected significance (BF) threshold to correct for multiple testing because of the large number of genetic markers (SNVs). To further correct for false positive QTL because of the large number of transcript expression traits, we computed a permutation-based False Discovery Rate (FDR) at 5%. We randomly selected 200 traits from our input phenotype file and permuted each of them 200 times. These 40,000 permuted phenotypes were used as input to call QTL using *cegwas2-nf* with EIGEN and BF threshold, respectively, as previously described^47, 49, 52^. Briefly, we used *BCFtools*^72^ to filter variants that had any missing genotype calls and variants that were below the 5% minor allele frequency. Then, we used *-indep-pairwise 50 10 0.8* in *PLINK* v1.9^85, 86^ to prune the genotypes to 20,991 markers with a linkage disequilibrium (LD) threshold of *r*^2^ < 0.8 and then generated the kinship matrix using the *A.mat()* function in the R package *rrBLUP* (v4.6.1)^87^. The number of independent tests (*N_test_*) within the genotype matrix was estimated using the R package *RSpectra* (v0.16.0) (https://github.com/yixuan/RSpectra) and *correlateR* (0.1) (https://github.com/AEBilgrau/correlateR). The eigen- decomposition significance (EIGEN) threshold was calculated as -log_10_(0.05/*N_test_*). We used the *GWAS()* function in the *rrBLUP* package to perform the genome-wide mapping with the EMMA algorithm^32^. QTL were defined by at least one marker that was above the EIGEN or BF threshold. The EIGEN and BF %5 FDR was calculated as the 95 percentile of the significance of all the detected QTL under each threshold. The EIGEN and BF 5% FDR thresholds were 6.11 and 7.76, respectively.

#### eQTL mapping

We performed GWA mapping on the expression traits of the 25,849 transcripts as for permuted expression traits but using the EIGEN 5% FDR (6.11) as the threshold. We identified QTL with significance that also passed the Bonferroni 5% FDR threshold to locate the best estimate of QTL positions with the highest significance. We used the generic function *cor()* in R and Pearson correlation coefficient to calculate the phenotypic variance explained by each QTL. We used the *LD()* function from the R package *genetics* (v1.3.8.1.2) (https://cran.r-project.org/package=genetics) to calculate the LD correlation coefficient *r*^2^ among QTL for traits with multiple eQTL.

#### eQTL classification

Local eQTL were classified if the QTL was within a 2 Mb region surrounding the transcript. All other QTL were classified as distant.

### Heritability calculation

Heritability estimates were calculated for each of the 25,849 traits used for eQTL mapping as previously described^88^. Narrow-sense heritability (*h*^2^) was calculated with the phenotype file and pruned genotypes in eQTL mapping using the functions *mmer()* and *pin()* in the R package *sommer* (v4.1.2)^89^. Broad-sense heritability (*H*^2^) was calculated using expression of replicates of each strain and the *lmer* function in the R package *lme4* (v1.1.21) with the model phenotype ∼ 1 + (1|strain)^90^.

### Hotspot identification

We first filtered out distant eQTL in common hyper-divergent genomic regions of wild C. elegans strains. Common hyper-divergent regions were defined among our 206 strains (reference N2 excluded) as described previously^27^. Briefly, we divided the genome into 1 kb bins and calculated the percentage of 206 strains that are classified as hyper-divergent in each bin. Common hyper-divergent regions were defined as bins ≥ 5%^27^.

Distant eQTL hotspots were identified by dividing the genome into 0.5 cM bins and counting the number of non-divergent distant eQTL that mapped to each bin. Significance was determined as bins with more eQTL than the 99th percentile of a Poisson distribution using the maximum likelihood method and the function eqpois() in the R package EnvStats^1, 3, 9, 91^.

### Reanalysis of RIAILs eQTL

We reclassified eQTL detected in a previous study using microarray expression data from synchronized young adult populations of 208 recombinant inbred advanced intercross lines (RIAILs) derived from N2 and CB4856^9, 36^. A total of 2,540 eQTL from 2,196 probes were identified using linkage mappings^9^. We selected 2,387 eQTL of 2,054 probes that are in 2,003 live genes based on the probe-gene list in the R package *linkagemapping* (https://github.com/AndersenLab/linkagemapping) and the GTF file (WS276)^53^. We classified 1,208 local eQTL and 1,179 distant eQTL as described above. We further identified hotspots as above for 1,124 distant eQTL that are not in the hyper-divergent regions of CB4856.

### Population genetics

We use 851,105 biallelic SNVs with no missing calls among the 207 strains from the hard-filtered VCF 20200815 CeNDR release to calculate population genomic statistics. Tajima’s D, Watterson’s θ, and pi were all calculated using *scikit-allel*^92^. Each of these statistics was calculated for non-overlapping 1,000-bp windows across the genome.

### Fine mapping for causal genes underlying hotspots

For transcript expression traits with distant eQTL in hotspots, we performed fine mapping using the pipeline *cegwas2-nf* as previously described^47^. Briefly, we defined QTL regions of interest from the GWA mapping as +/- 100 SNVs from the rightmost and leftmost markers above the EIGEN 5% FDR significance threshold. Then, using genotype data from the imputed hard-filtered isotype VCF (20200815 CeNDR release), we generated a QTL region of interest genotype matrix that was filtered as described above, with the one exception that we did not perform LD pruning. We used *PLINK* v1.9^85, 86^ to extract the LD between the markers used for fine mapping and the QTL peak marker identified from GWA mappings. We used the same command as above to perform fine mappings. To identify causal genes and variants that affect expression of several transcripts underlying hotspots, we retained the fine-mapped candidate variants that passed the following per trait filters: top 5% most significant markers; out of common hyper-divergent genomic regions; with negative BLOSUM^93^ scores as characterized and annotated in CeNDR^25^.

### Enrichment analysis

Gene set enrichment analyses were carried out for all genes found with transcript-level eQTL and for genes with distant eQTL in each hotspot using the web-based tool *WormCat*^39^.

### Mediation analysis

#### GWA mapping of diverse C. elegans phenotypes

We obtained nine different phenotype data used in previous *C.elegans* natural variation and GWA studies^43, 44, 46–52^. We filtered genetically distinct isotype strains for each trait based on CeNDR (20200815 release) and performed GWA mapping as for permuted expression traits but mostly using EIGEN or BF as the threshold according to the original studies. GWA was performed under EIGEN for two studies originally using BF as the threshold^48, 49^.

#### Mediation analysis

For each QTL of the above phenotypes, we used the genotype (*Exposure*) at the phenotype QTL peak, transcript expression traits (*Mediator*) that have eQTL overlapped with the phenotype QTL, and the phenotype (*Outcome*) as input to perform mediation analysis using the *medTest()* function and 1,000 permutations for *p*-value correction in the R package *MultiMed* (v2.6.0) (https://bioconductor.org/packages/release/bioc/html/MultiMed.html). For mediation, we used only strains with all of the three input data types available and where variation was found. For instance, between the 202 strains used in the study of ABZ resistance^44^ and the 207 strains used in this study, 167 strains overlapped. Although we searched overlapped eQTL against QTL in the GWA mapping for ABZ resistance using 202 strains (Supplementary Fig. 8), 167 strains at maximum were used in mediation analysis. Furthermore, because some transcripts were found in hyper-divergent regions in certain strains and their expression data were filtered out, the rest of the strains with all of the data types available might contain no variation in one or all of the three data types and were not used in mediation analysis. For example, we found 1,193 eQTL overlapped with the QTL on chromosome II in the GWA mapping for ABZ resistance, but only 1,157 mediation analyses were performed.

For mediators with adjusted *p* < 0.05 or mediation estimate greater than the 99^th^ percentile of the distribution of mediation estimates, we performed a second mediation analysis as described previously^9^ using the *mediate()* function from the R package *mediation* (version 4.5.0)^94^ to filter out the uninterpretable results where the proportion of the total effect (the estimated effect of genotype on phenotype, ignoring expression) that can be explained by the mediation effect (the estimated effect of expression on phenotype) is negative or larger than 100%.

#### GWA of traits regressed by transcript expression

We regressed the trait animal length (q90.TOF)^44^ by expression of every transcript using the generic function *residuals()* in R, which fits a linear model with the formula (*phenotype* ∼ *expression*) to account for any differences in phenotype parameters present in transcript expression. Then GWA was performed for each regressed trait as for permuted expression traits using BF as the threshold.

## Acknowledgements

We would like to thank Stefan Zdraljevic and Samuel J. Widmayer for helpful suggestions. G.Z. is supported by the NSF-Simons Center for Quantitative Biology at Northwestern University (awards Simons Foundation/SFARI 597491-RWC and the National Science Foundation 1764421). S.R.H. was funded by a DFG fellowship (HA 8449/1-1) from the Deutsche Forschungsgemeinschaft (www.dfg.de). E.C.A. is supported by a National Science Foundation CAREER Award (IOS-1751035) and a grant from the National Institutes of Health R01 DK115690. The *C. elegans* Natural Diversity Resource is supported by a National Science Foundation Living Collections Award to R.E.T. and E.C.A. (1930382). We also like to thank WormBase for which these analyses would not have been possible.

## Author contributions

E.C.A. conceived of and designed the study. D.L., S.R.H., and G.Z. prepared *C. elegans* cultures. G.Z. and N.M.R. performed RNA-seq experiments. G.Z. analyzed the data. G.Z. and E.C.A. wrote the manuscript.

## Competing interests

The authors declare no competing interests.

## Data and code availability

The raw RNA-seq data generated in this study are available at the NCBI Sequence Read Archive (Project PRJNA669810). The raw expression counts and TPM quantified in this study are available at NCBI’s Gene Expression Omnibus (Series GSE186719). The RNA-seq mapping pipeline can be found at https://github.com/AndersenLab/PEmRNA-seq-nf. The mediation analysis pipeline can be found at https://github.com/AndersenLab/mediation_GWAeQTL. The datasets and code for generating all figures can be found at https://github.com/AndersenLab/WI-Ce-eQTL.

## Supplementary Information

### Description of Additional Supplementary Files

File Name: Supplementary Data 1

Description: Expression abundances and heritabilities of 25,849 transcripts in 207 wild C. elegans strains

File Name: Supplementary Data 2

Description: GWA mapping results of 5,291 transcript expression traits, eQTL classification, and distant eQTL hotspots

File Name: Supplementary Data 3

Description: Enrichment of genes with eQTL, distant eQTL in hotspots

File Name: Supplementary Data 4

Description: Fine mappings for distant eQTL in hotspots

### Supplementary Figures

**Supplementary Fig. 1.**
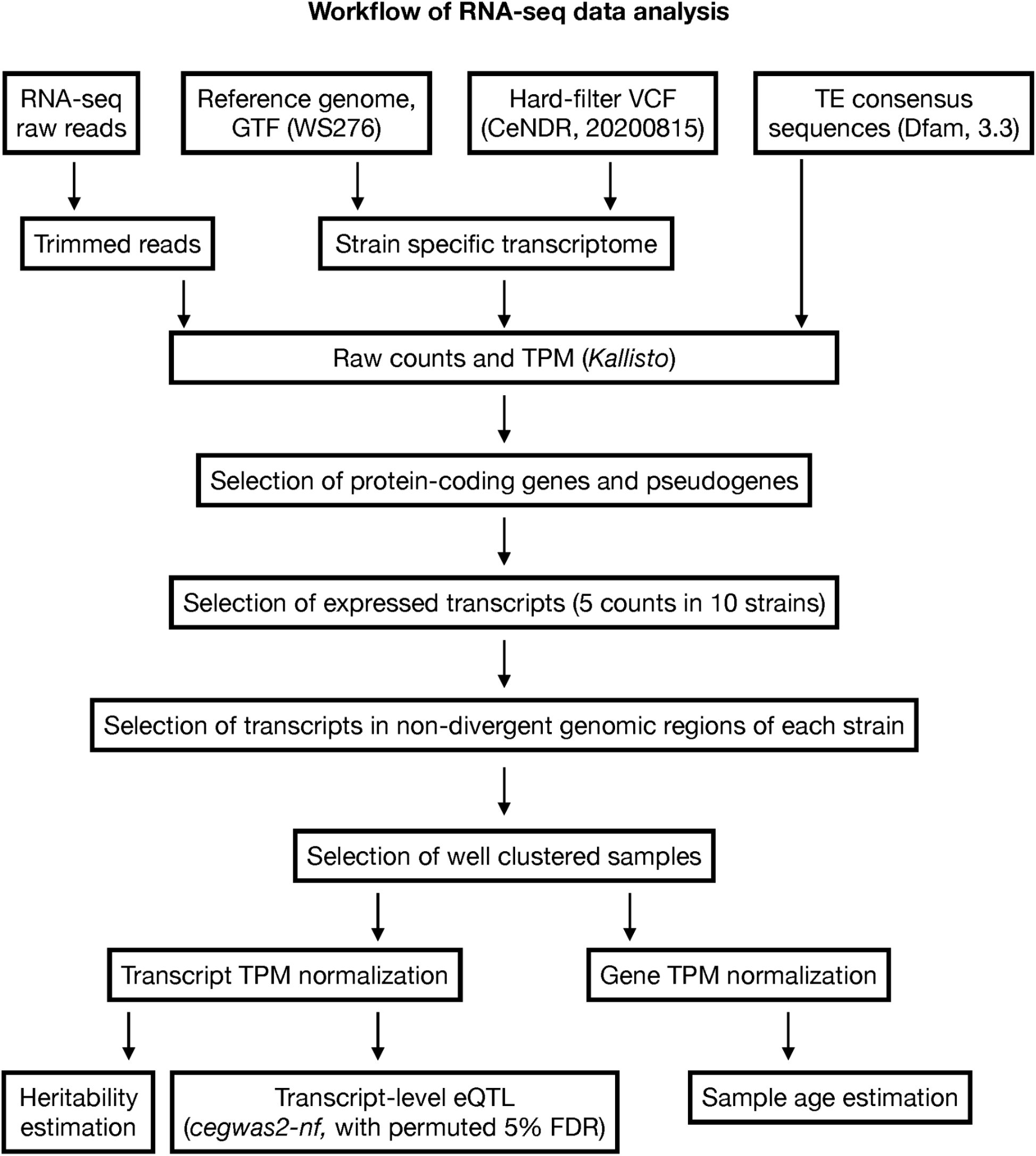
Workflow of RNA-seq data processing.

**Supplementary Fig. 2.**
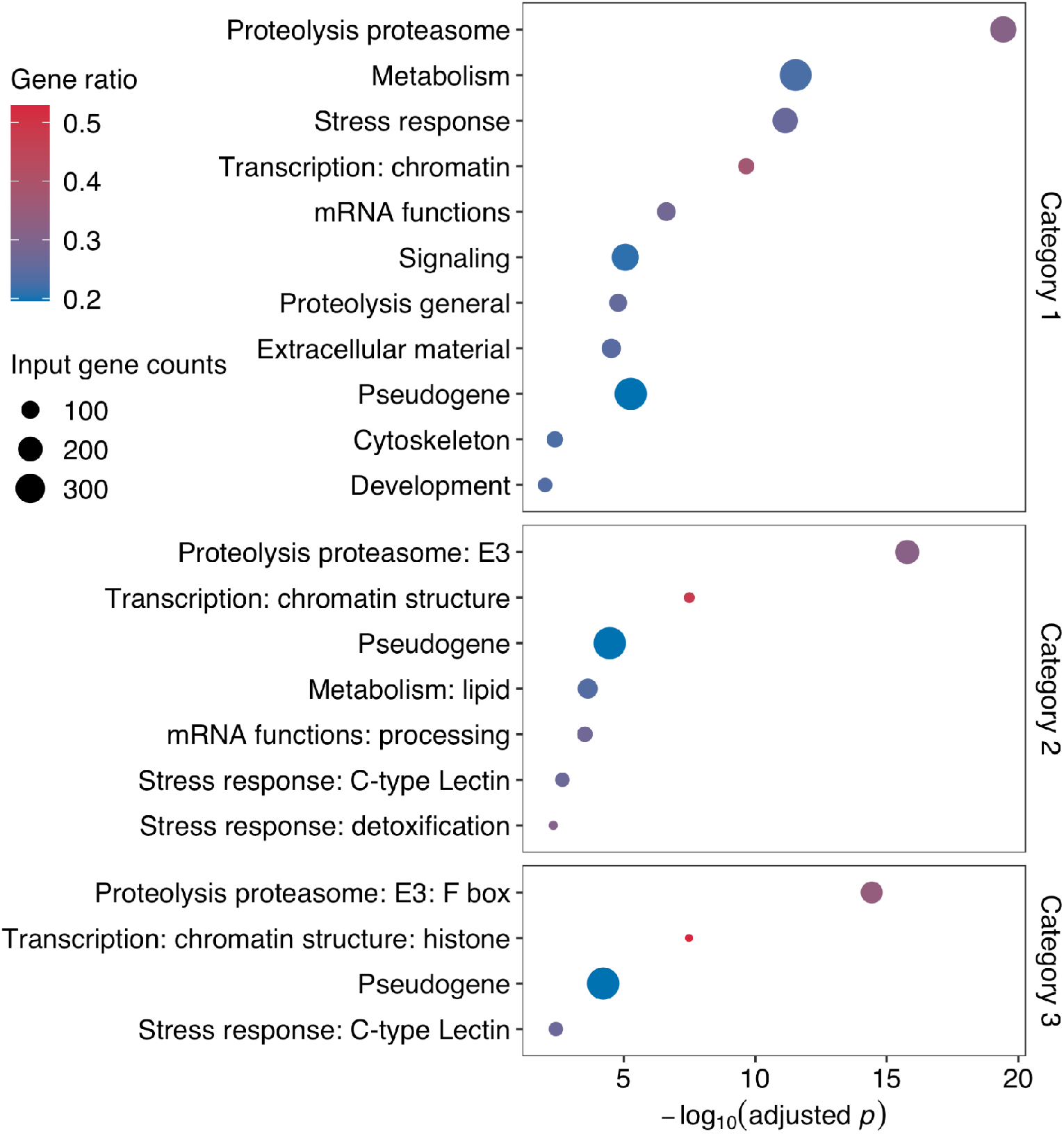
Gene set enrichment analysis for genes with transcript level eQTL. Enriched gene classes of broad and specific categories (Category 1 to 3)^39^ are shown on the y axis. Bonferroni FDR corrected significance values using Fisher Exact Test for gene set enrichment analysis are shown on the x axis. The sizes of the circles correspond to the input gene counts of the annotation and the colors of the circles correspond to the gene ratio of input gene counts to total gene counts of the annotation.

**Supplementary Fig. 3.**
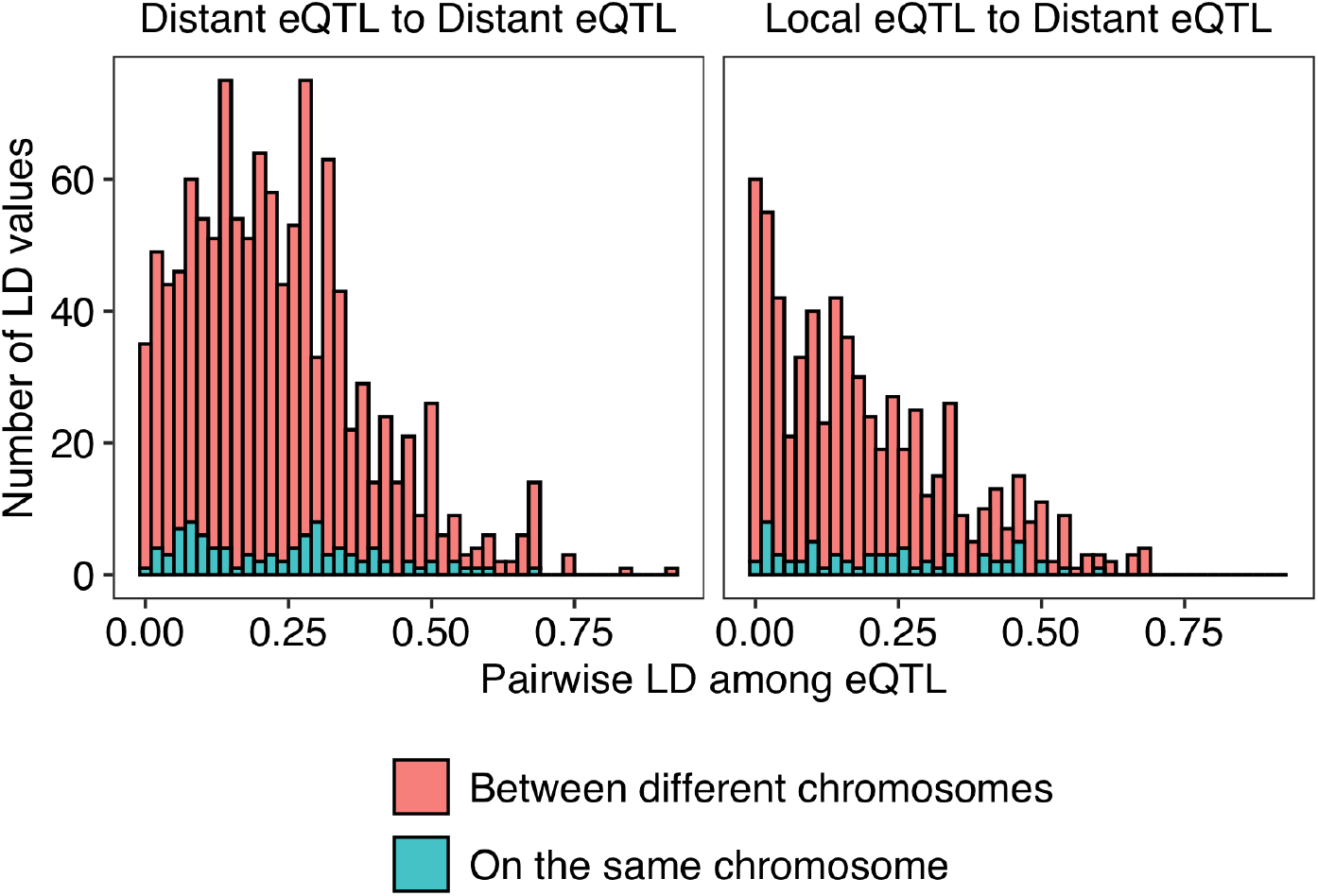
A histogram showing the distribution of linkage disequilibrium (LD) values (x-axis) among QTL of multiple eQTL of transcript expression traits. A total of 861 traits were found with multiple eQTL. LD of eQTL from the same chromosome and different chromosomes are colored salmon and blue, respectively.

**Supplementary Fig. 4.**
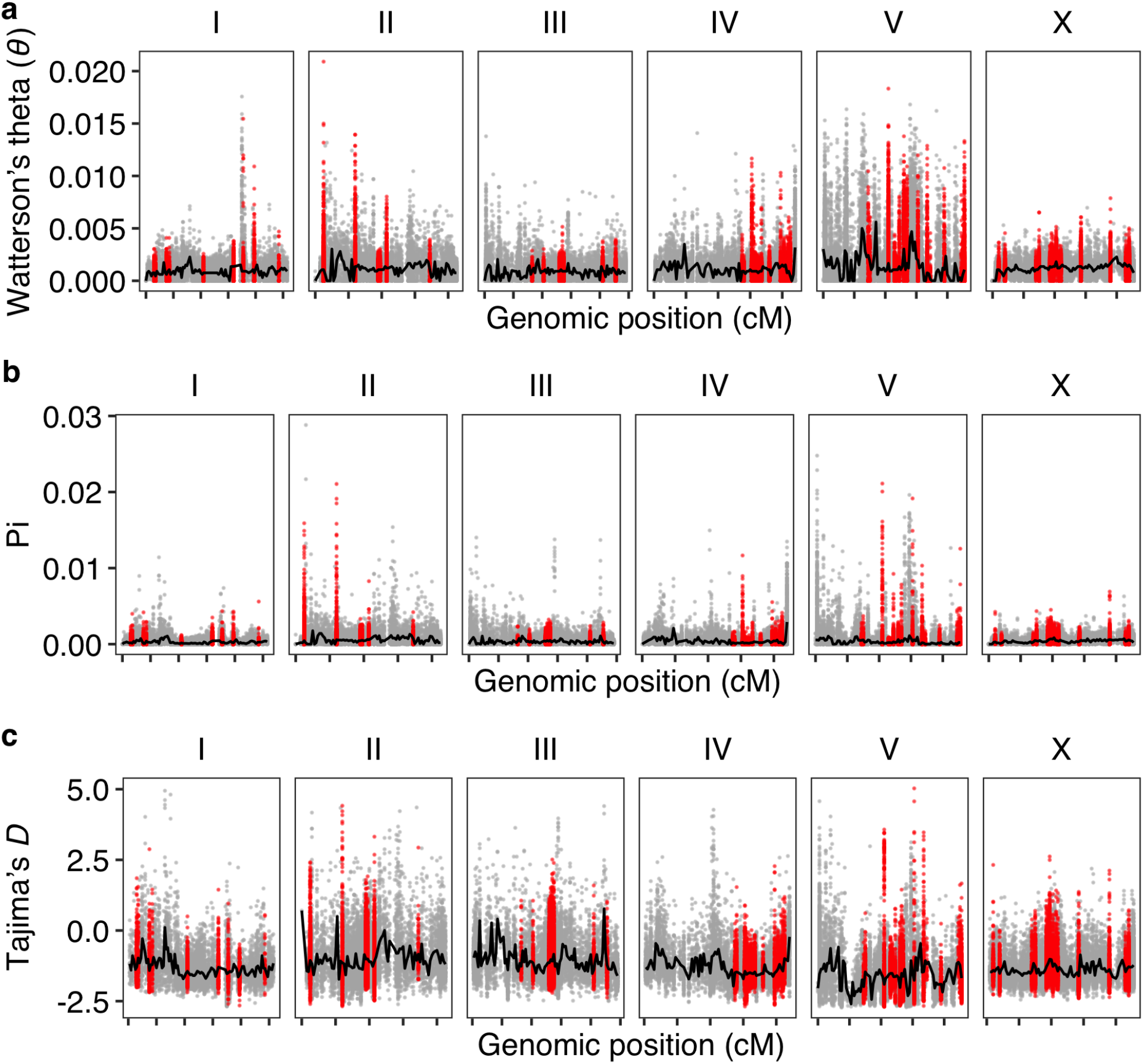
Genome-wide pattern of (a) Watterson’s theta (*θ*), (b) nucleotide diversity (pi), and (c) Tajima’s *D*. Each point represents the value (y-axis) for a non-overlapping 1 kb genomic window and is plotted against the genome position (x-axis) with tick marks denoting every 10 cM. Points for genomic windows in distant eQTL hotspots are colored red. Other points are colored gray. Median values of each statistic in each 0.5 cM bin were colored black. Tajima’s D values that suggest purifying selection are outliers for most values within a hotspot.

**Supplementary Fig. 5.**
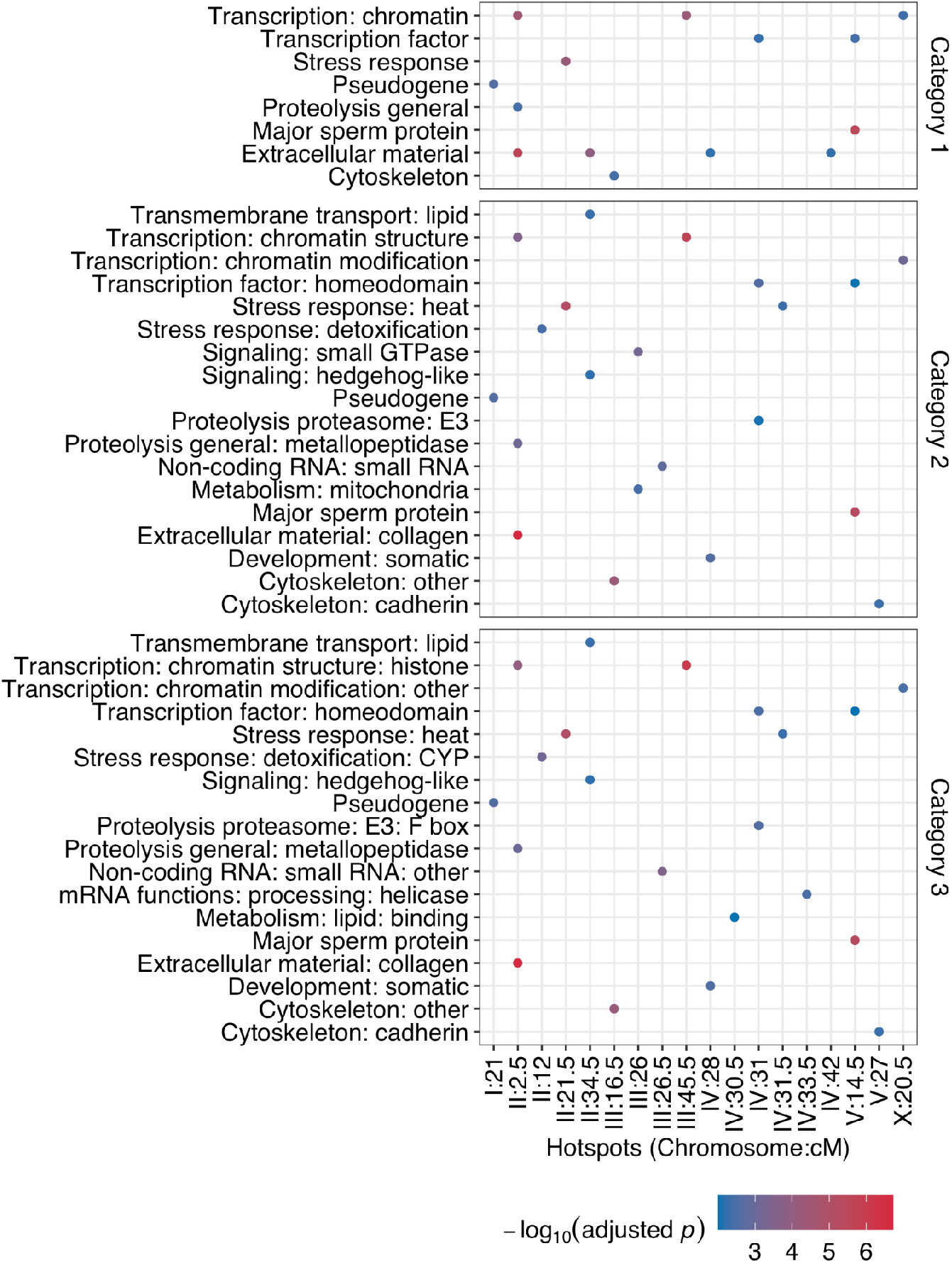
Gene set enrichment analysis for genes with transcript level distant eQTL in each hotspot. Broad and specific categories of enriched gene (Category 1 to 3)^39^ are shown on the y axis. Distant eQTL hotspots with significant gene set enrichment are shown on the x axis. The colors of the circles correspond to Bonferroni FDR corrected significance values using Fisher Exact Test.

**Supplementary Fig. 6.**
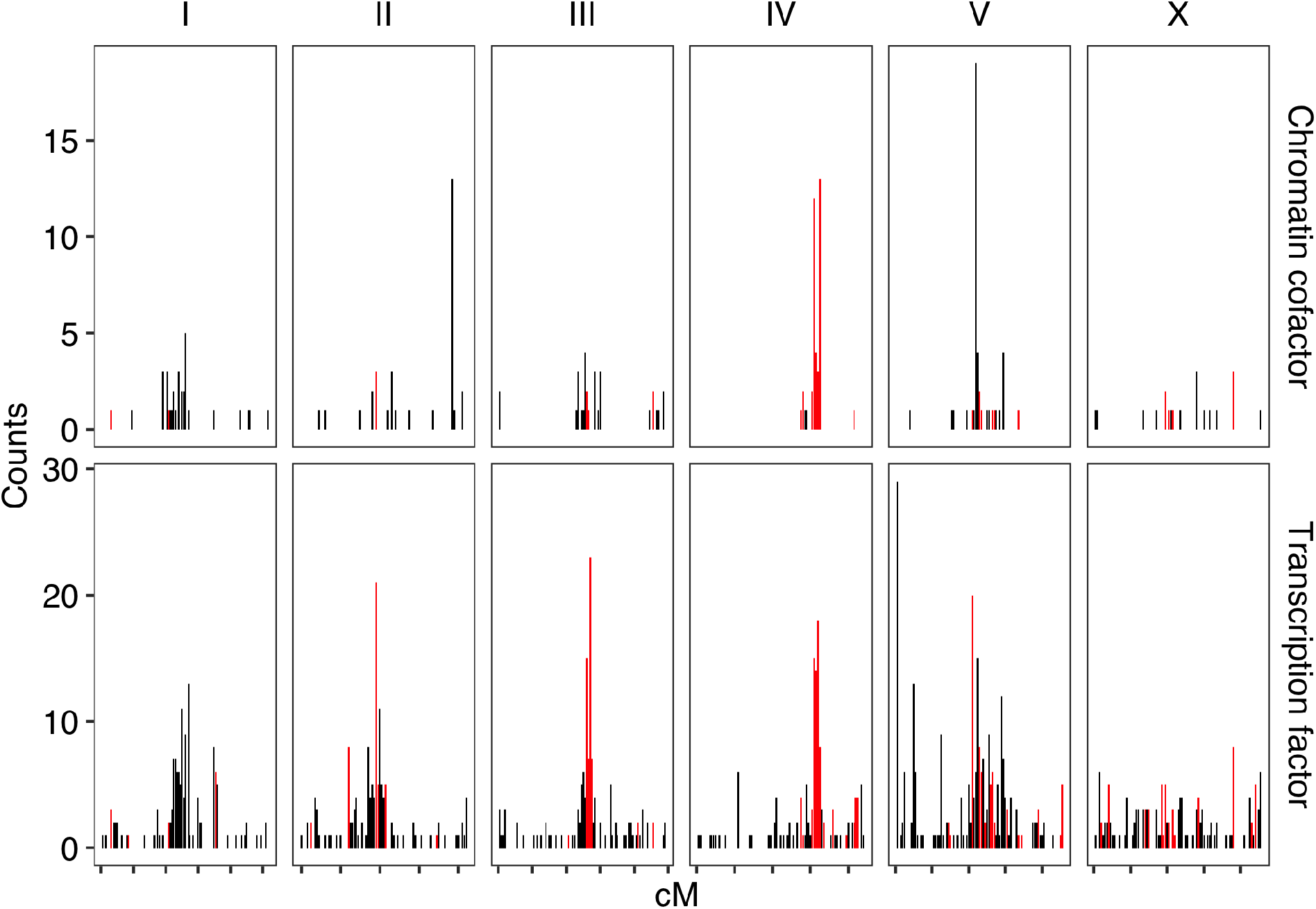
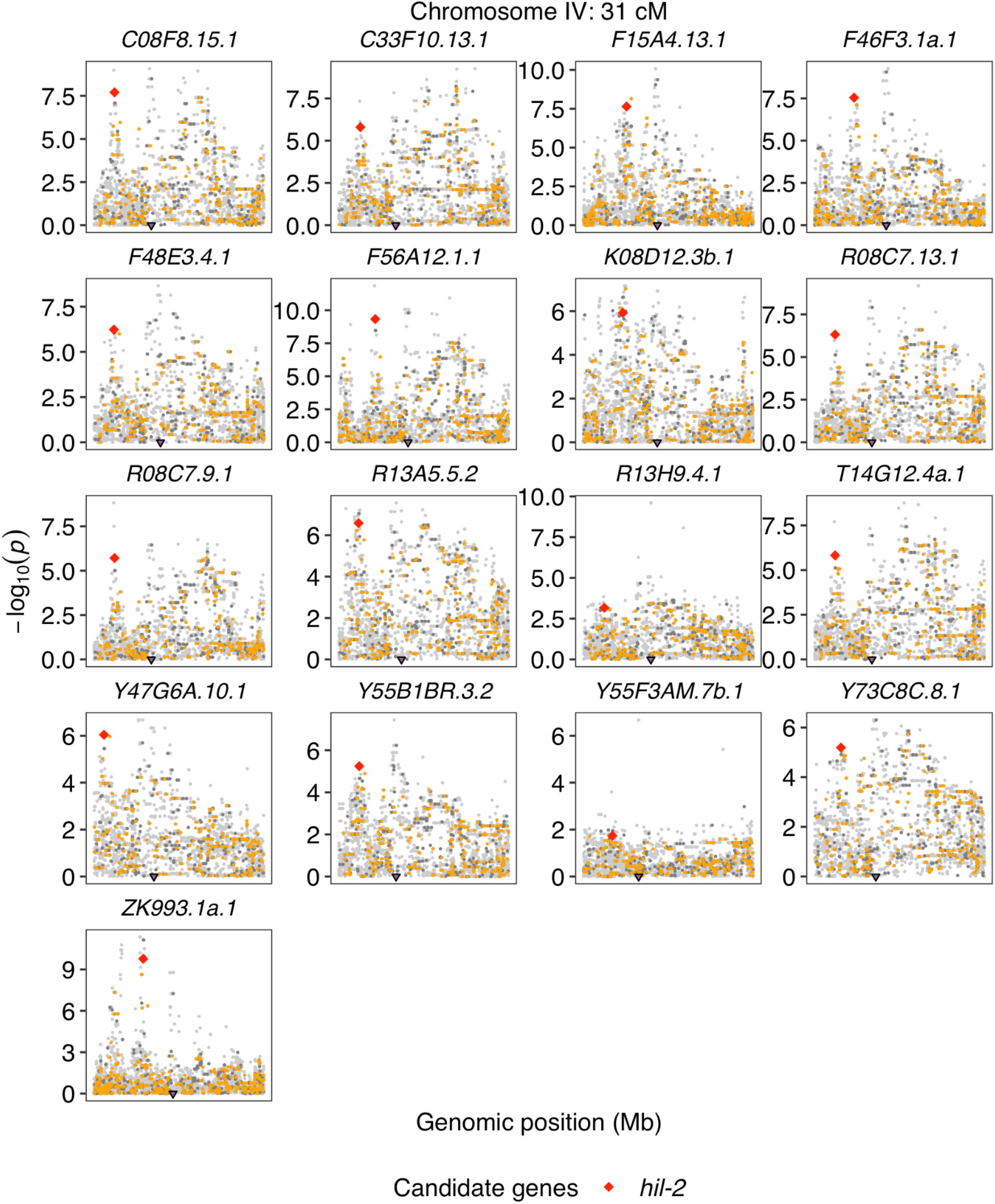
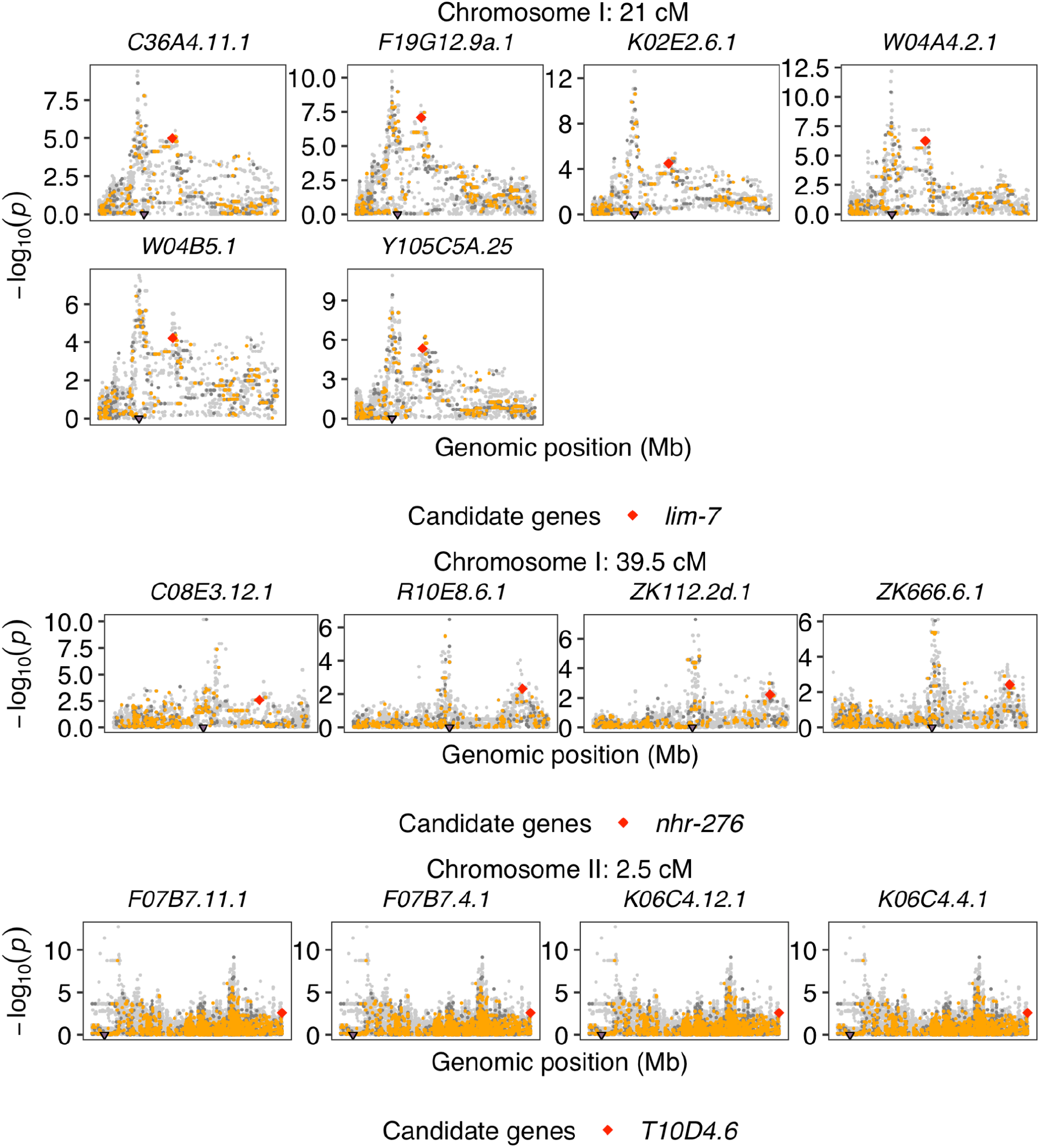
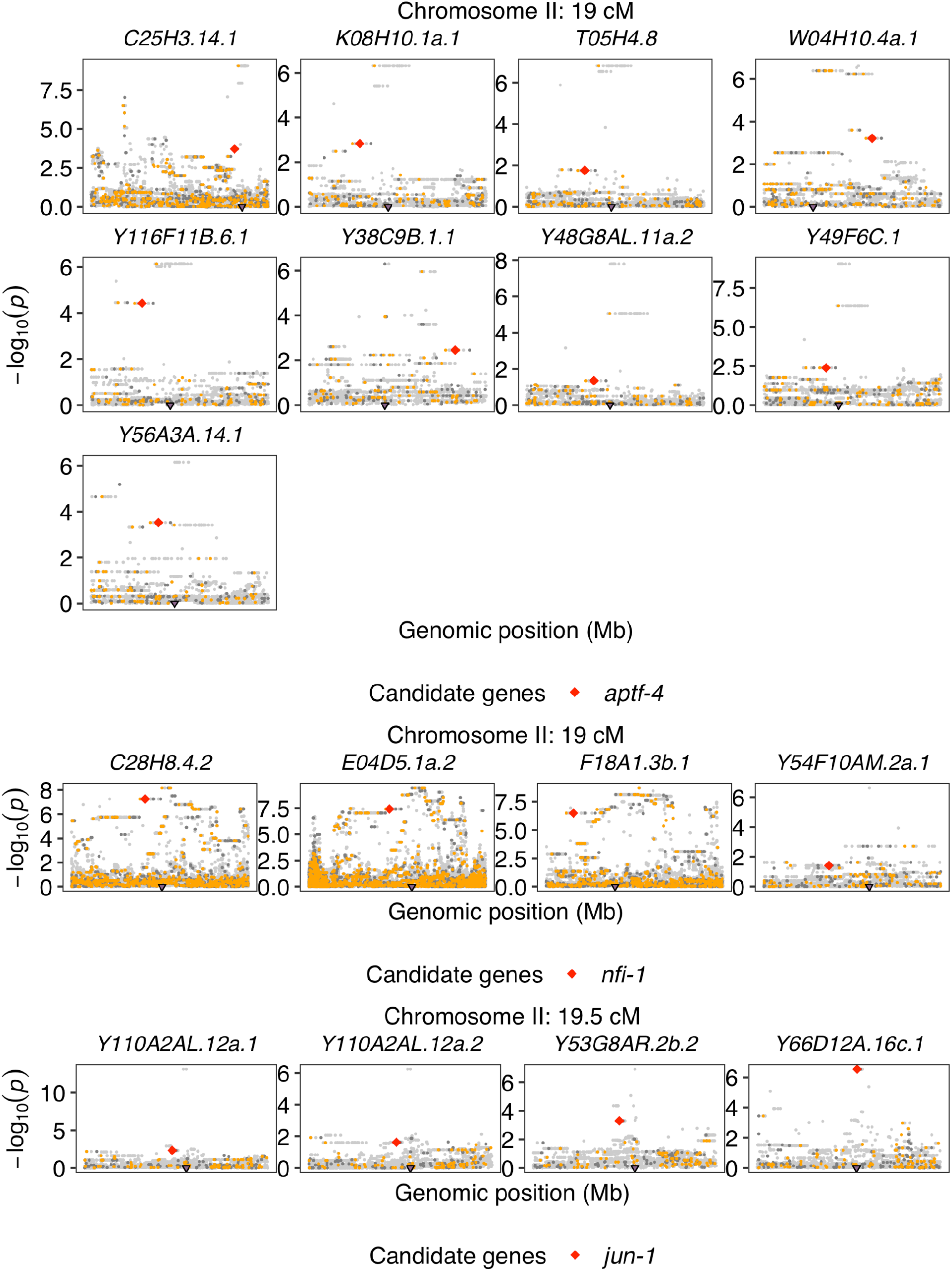
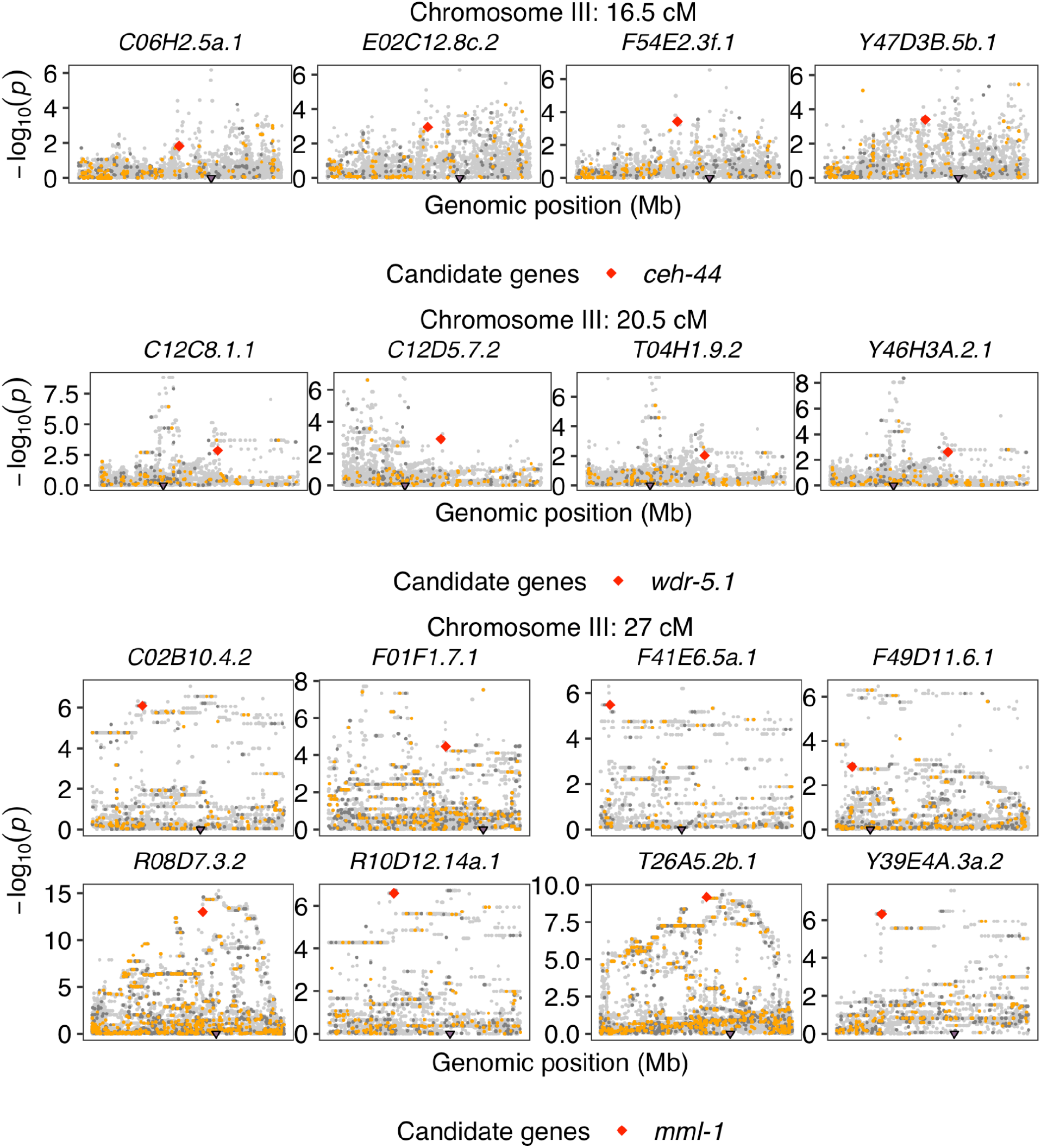
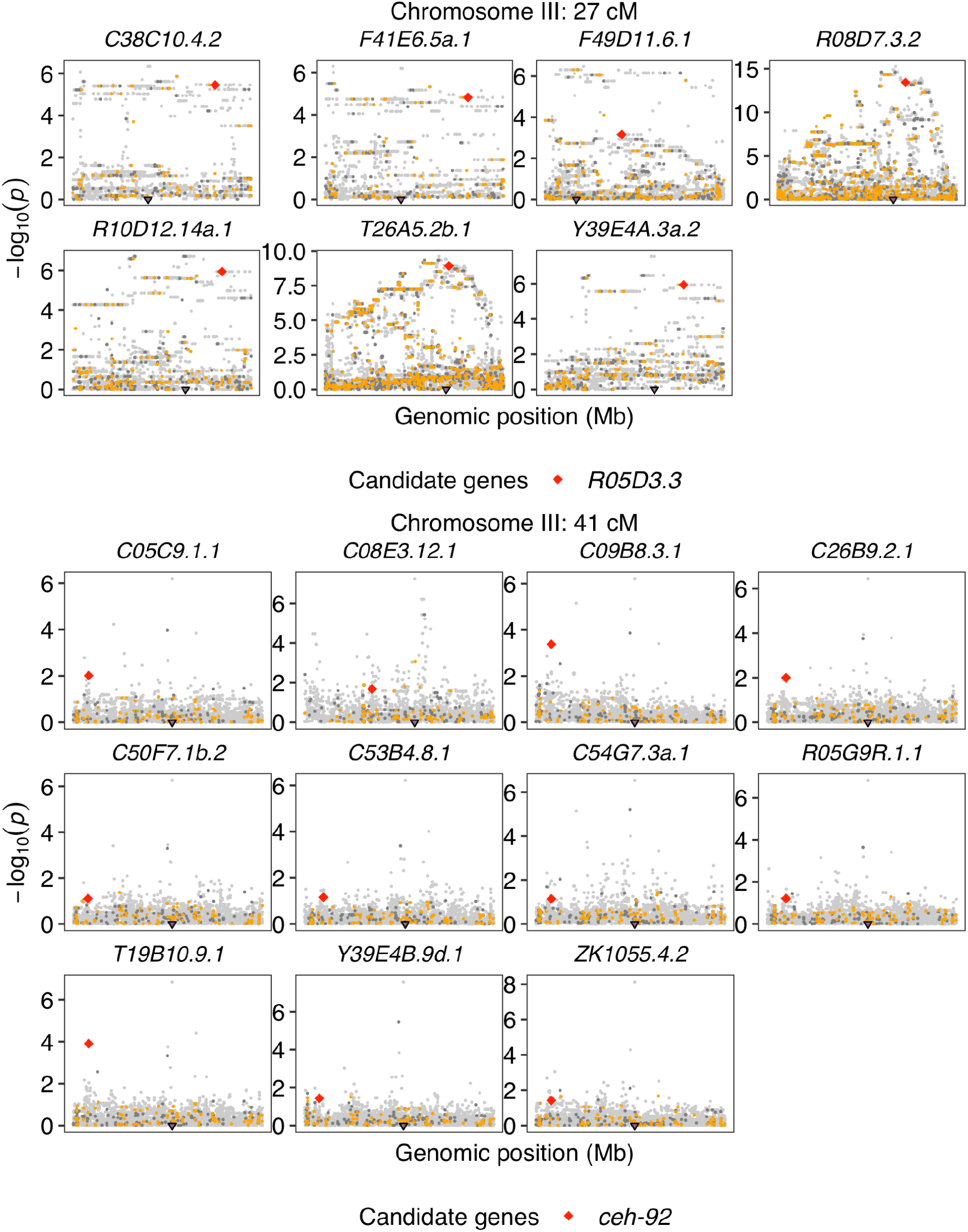
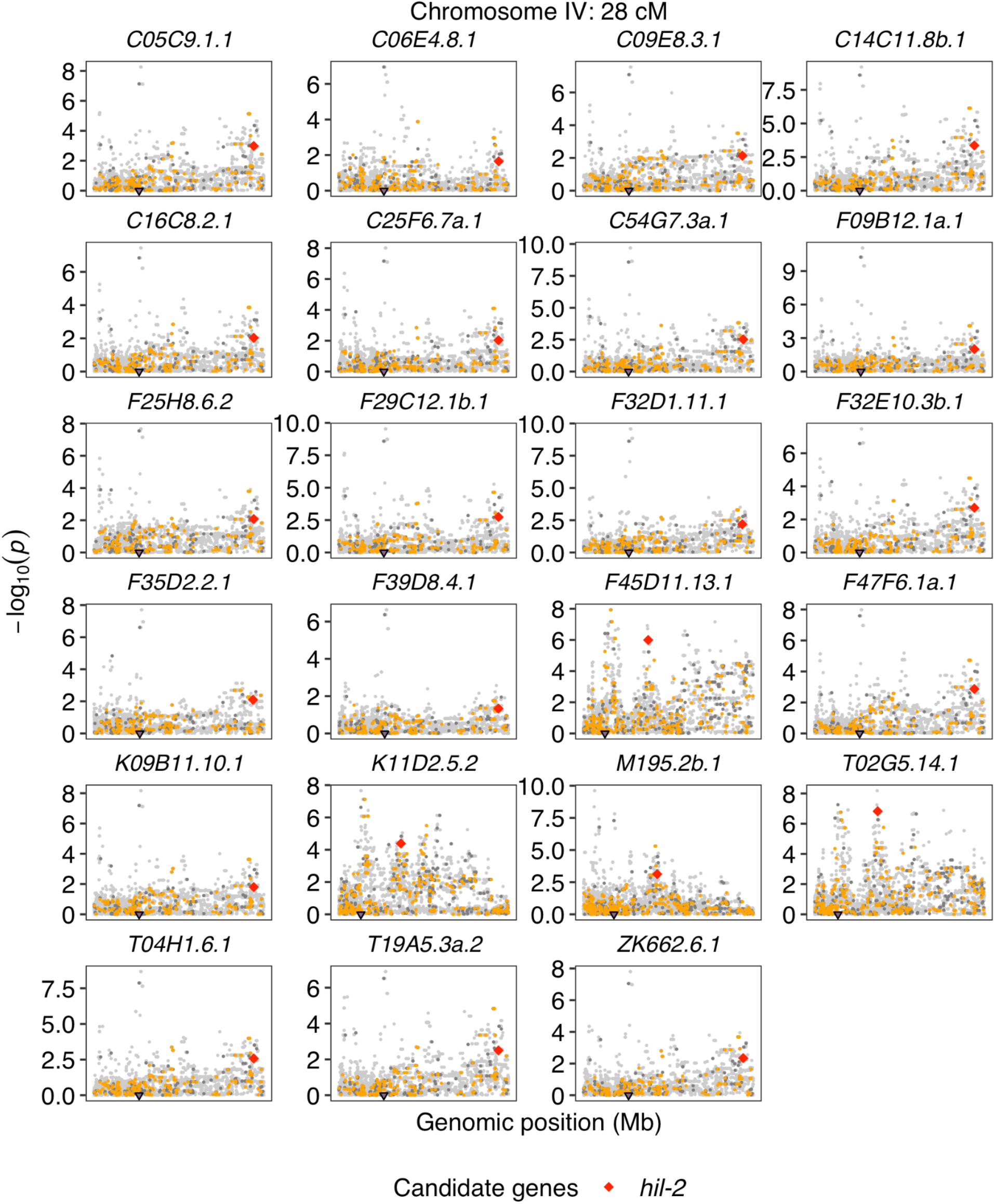
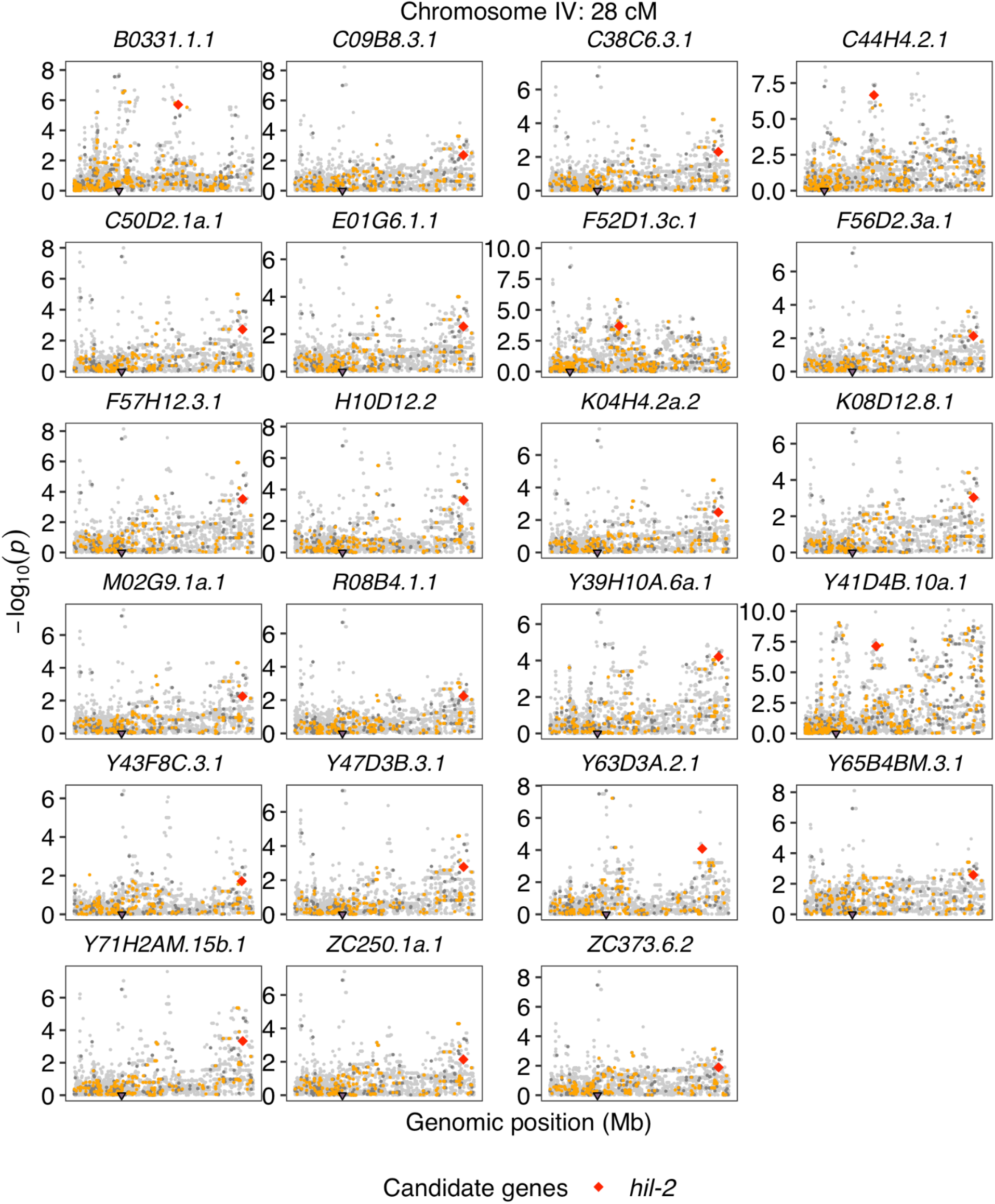
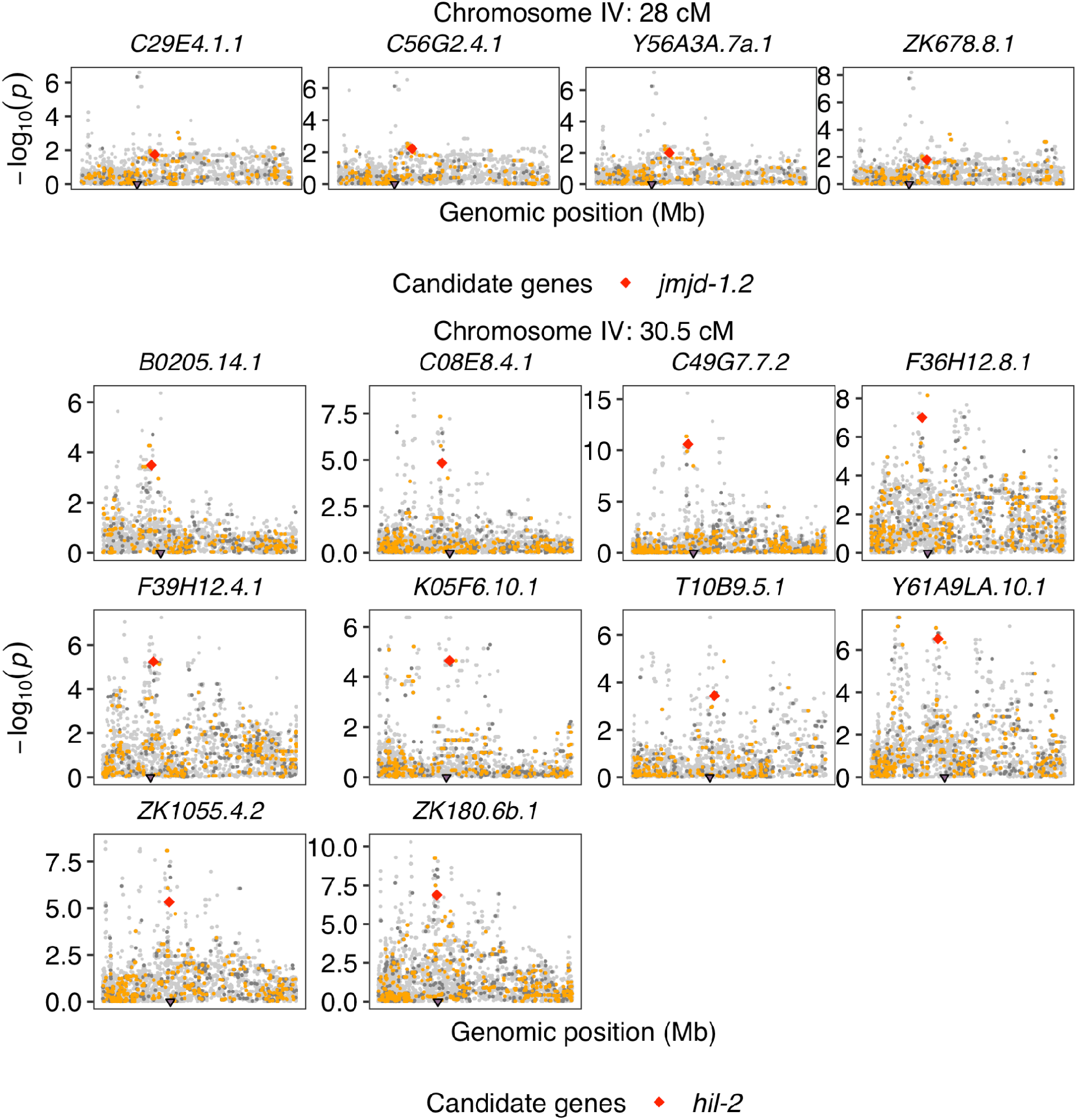
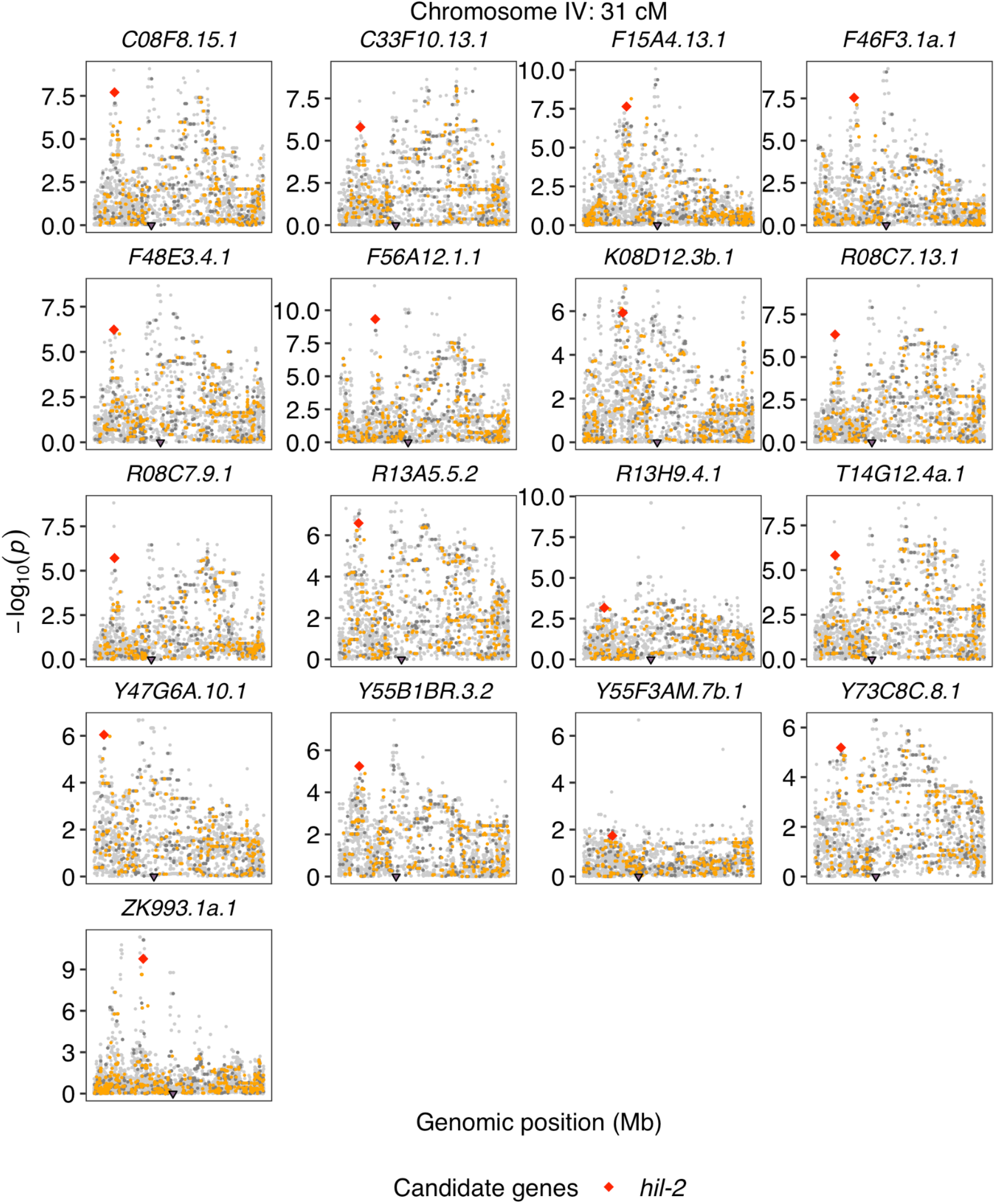
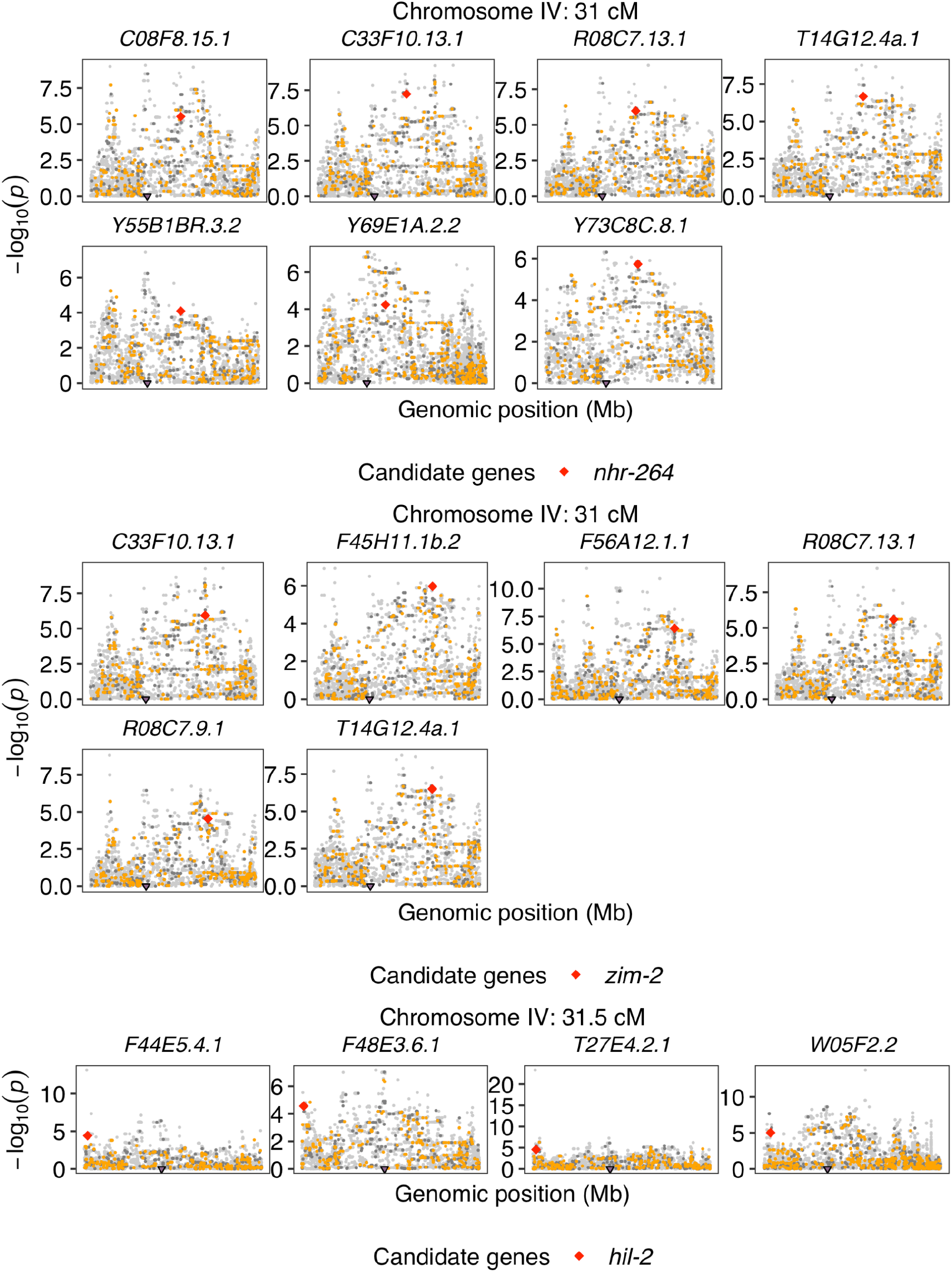
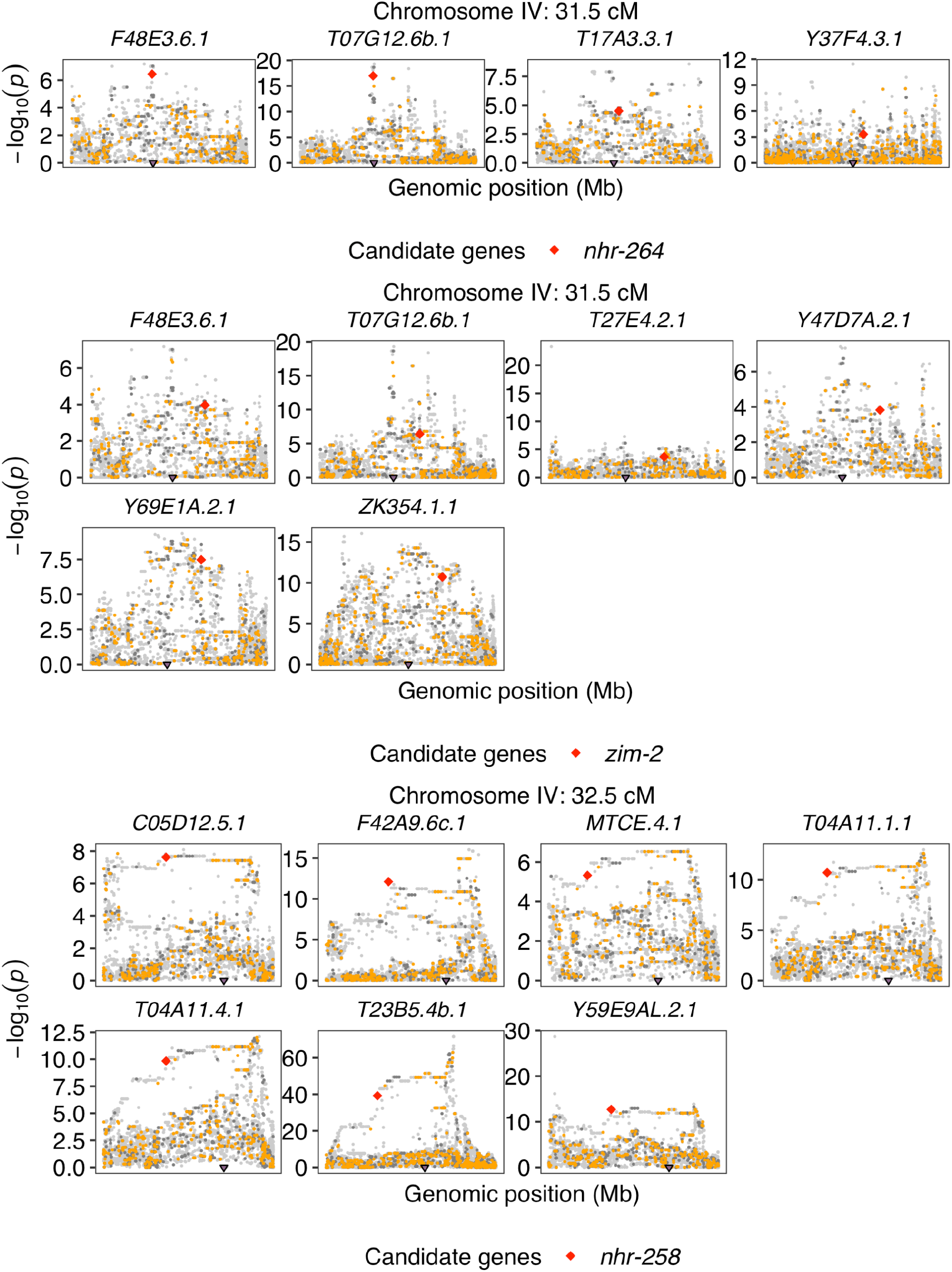
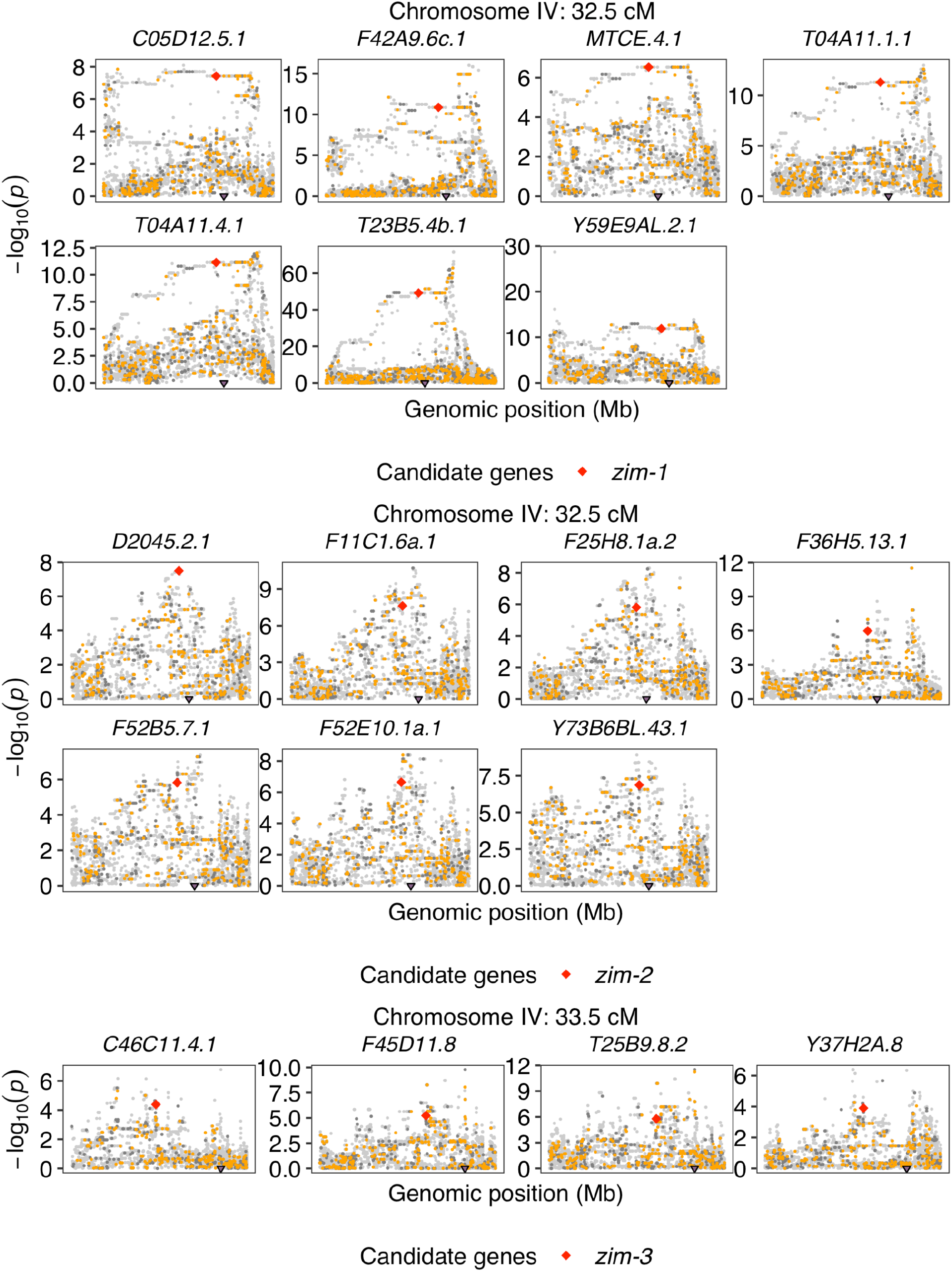
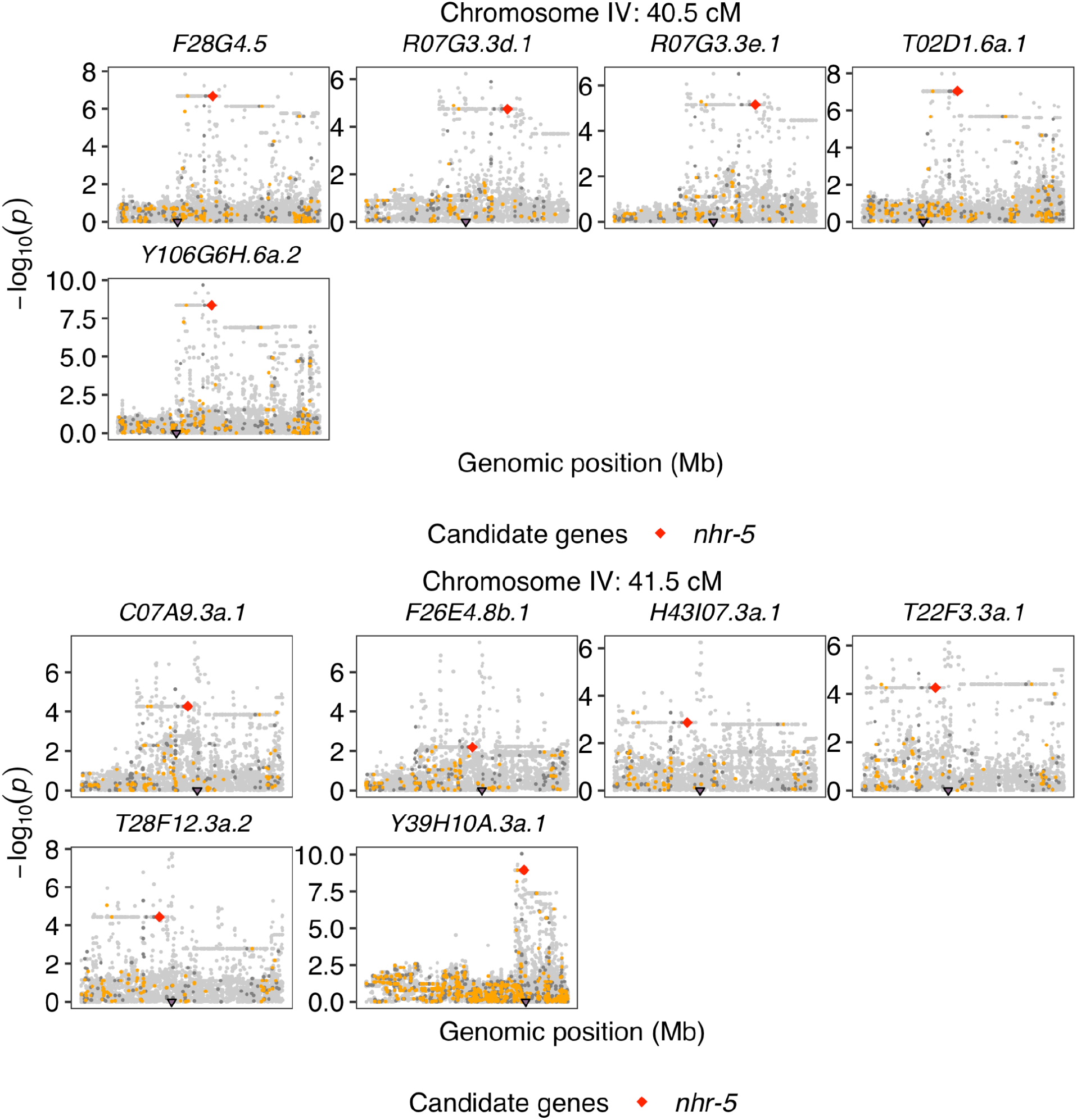
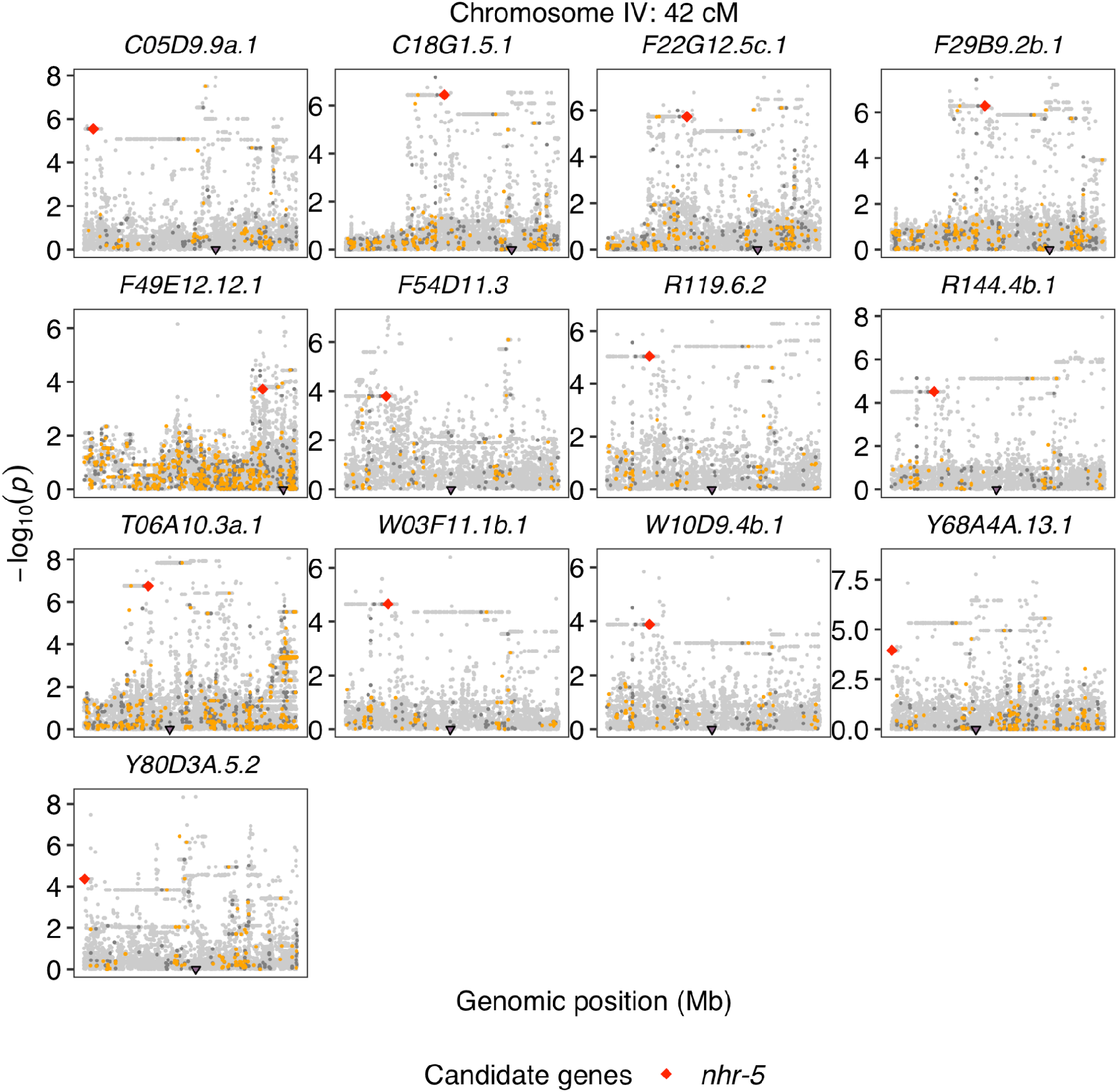
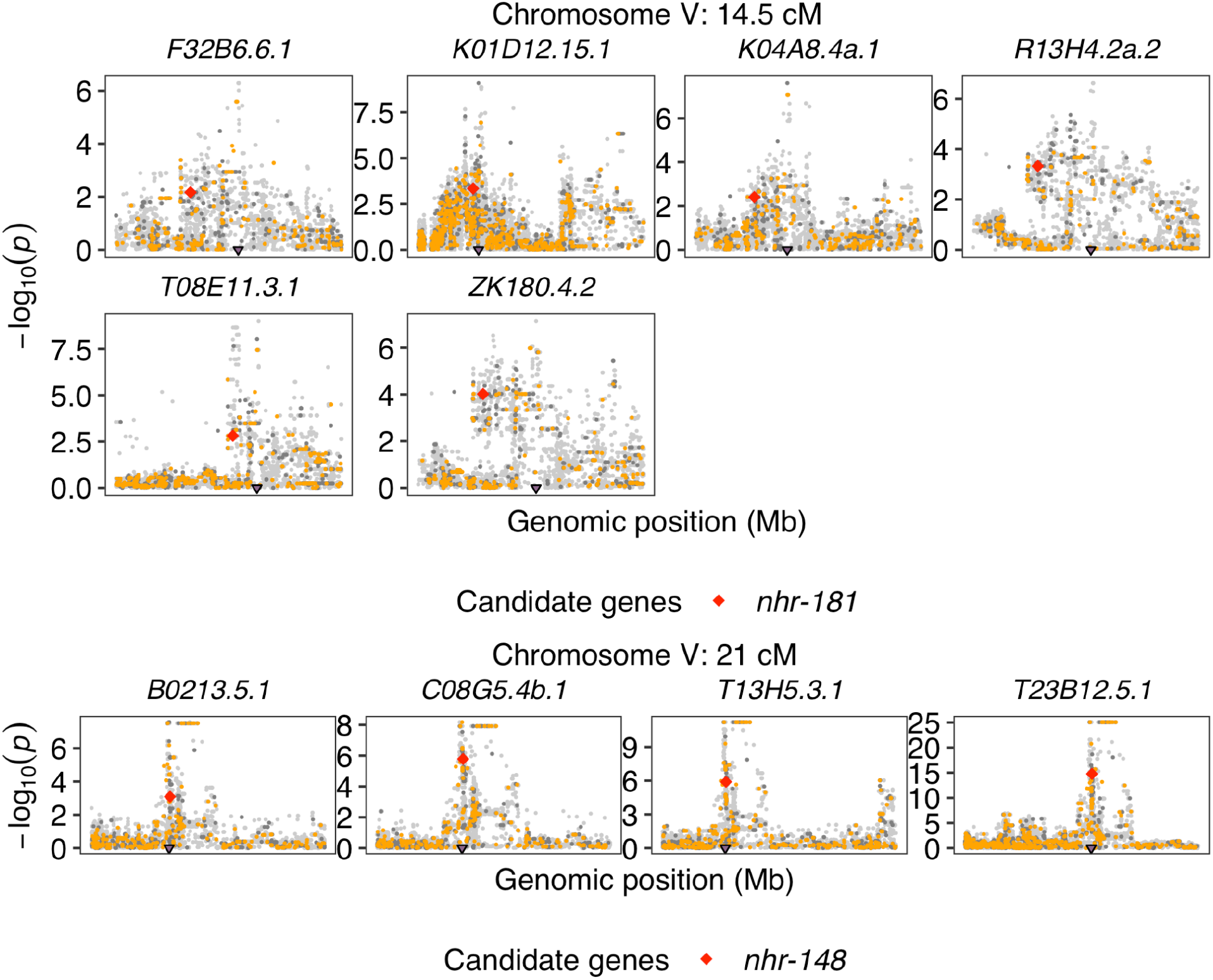
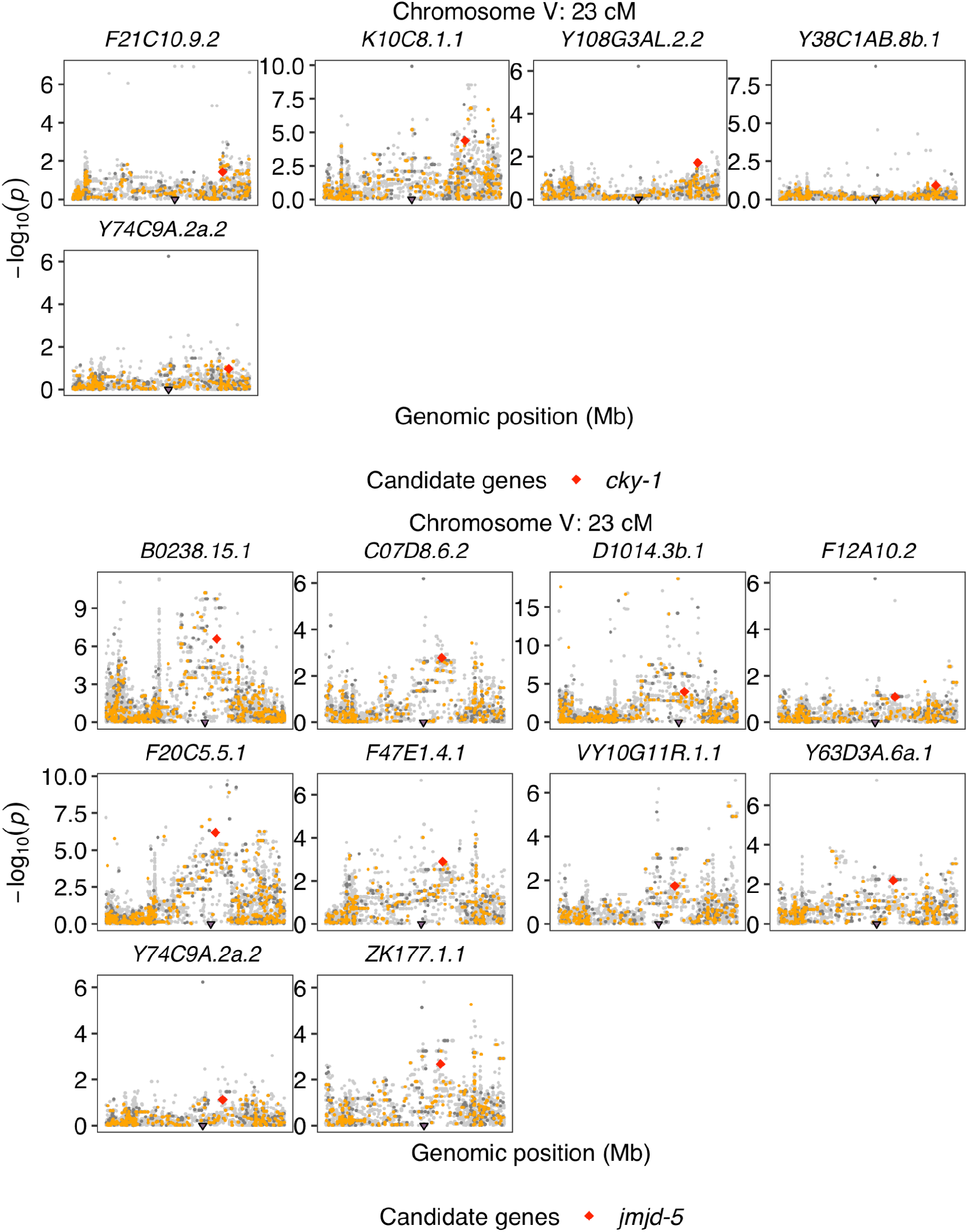
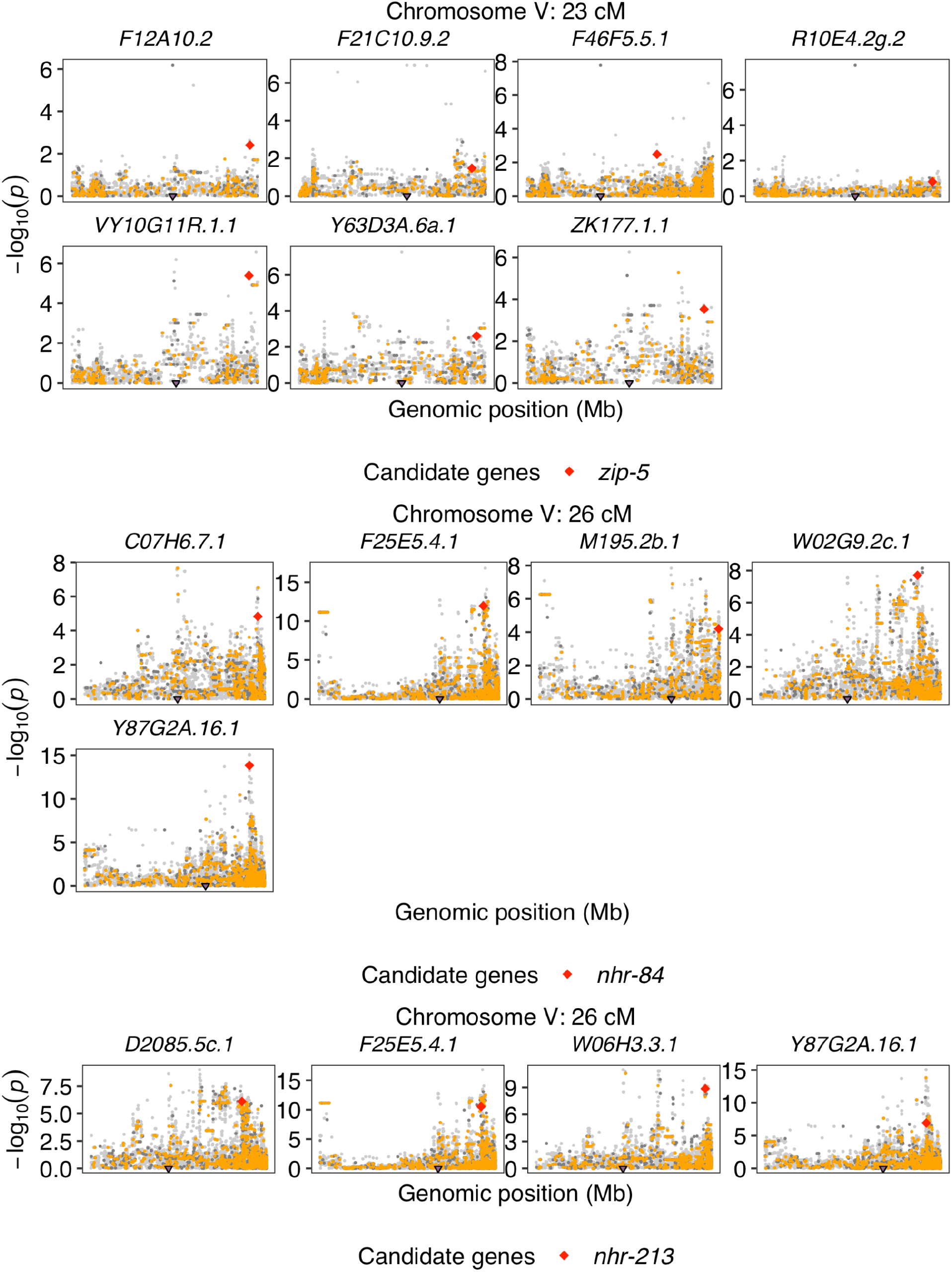
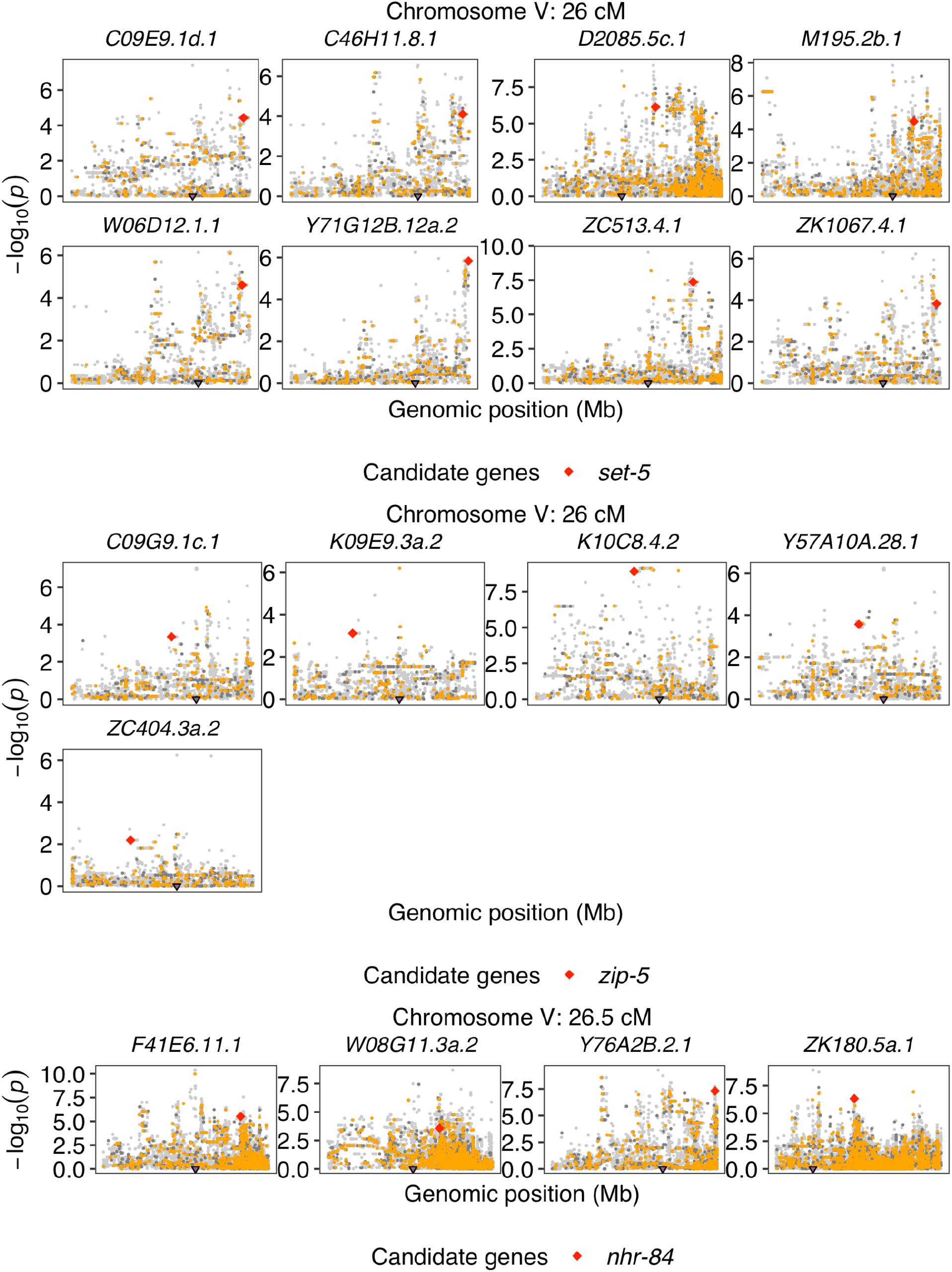
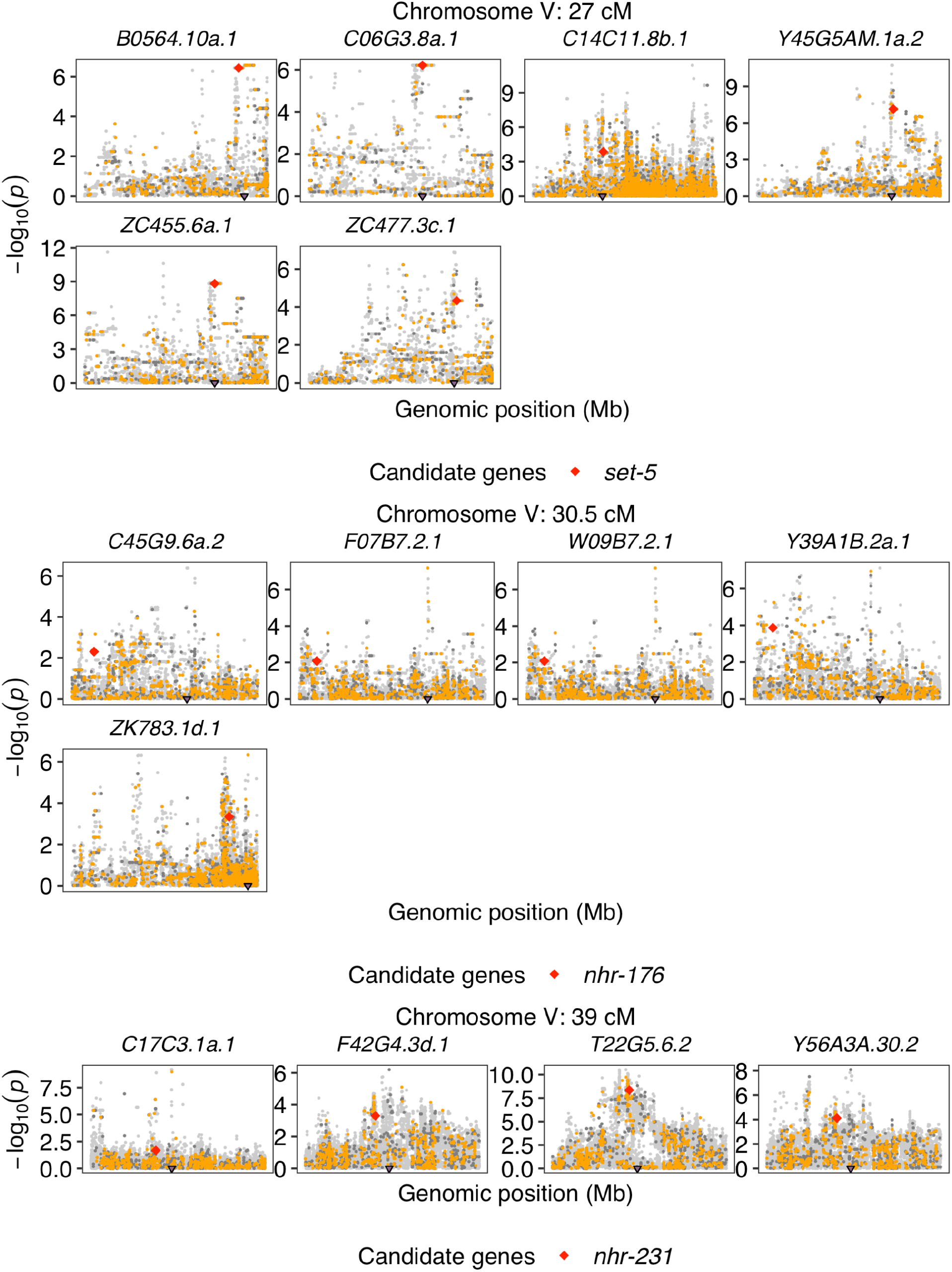
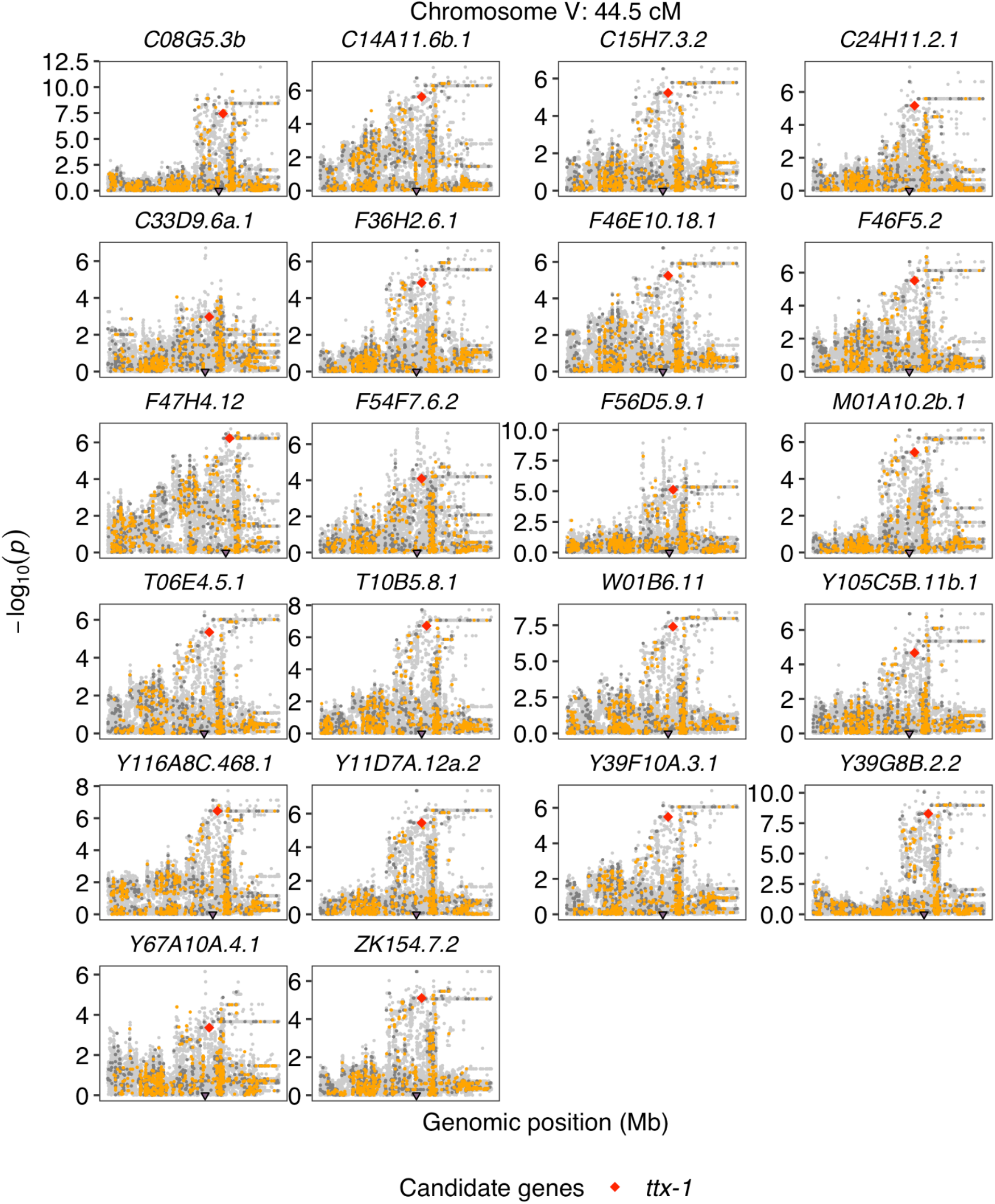
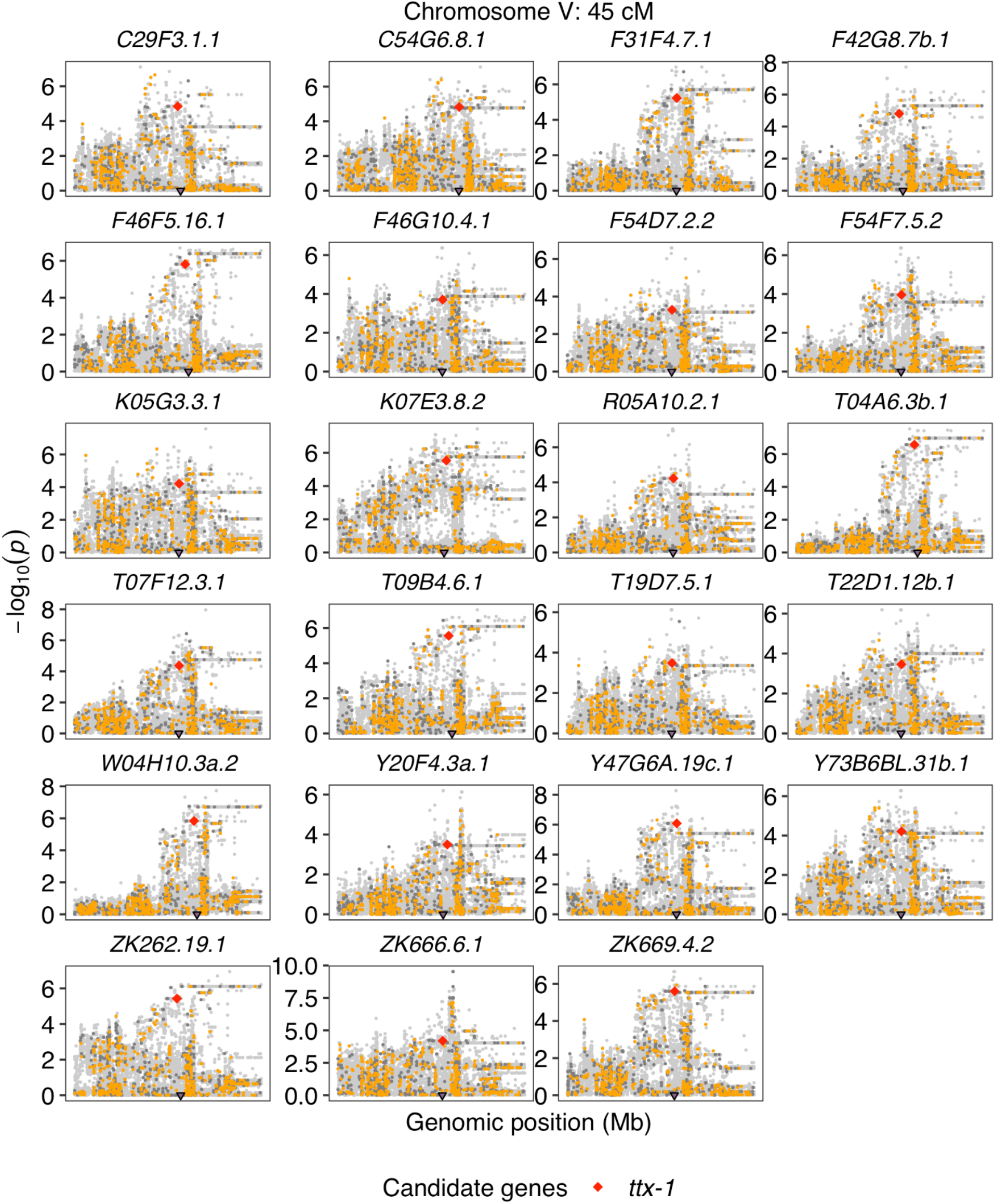
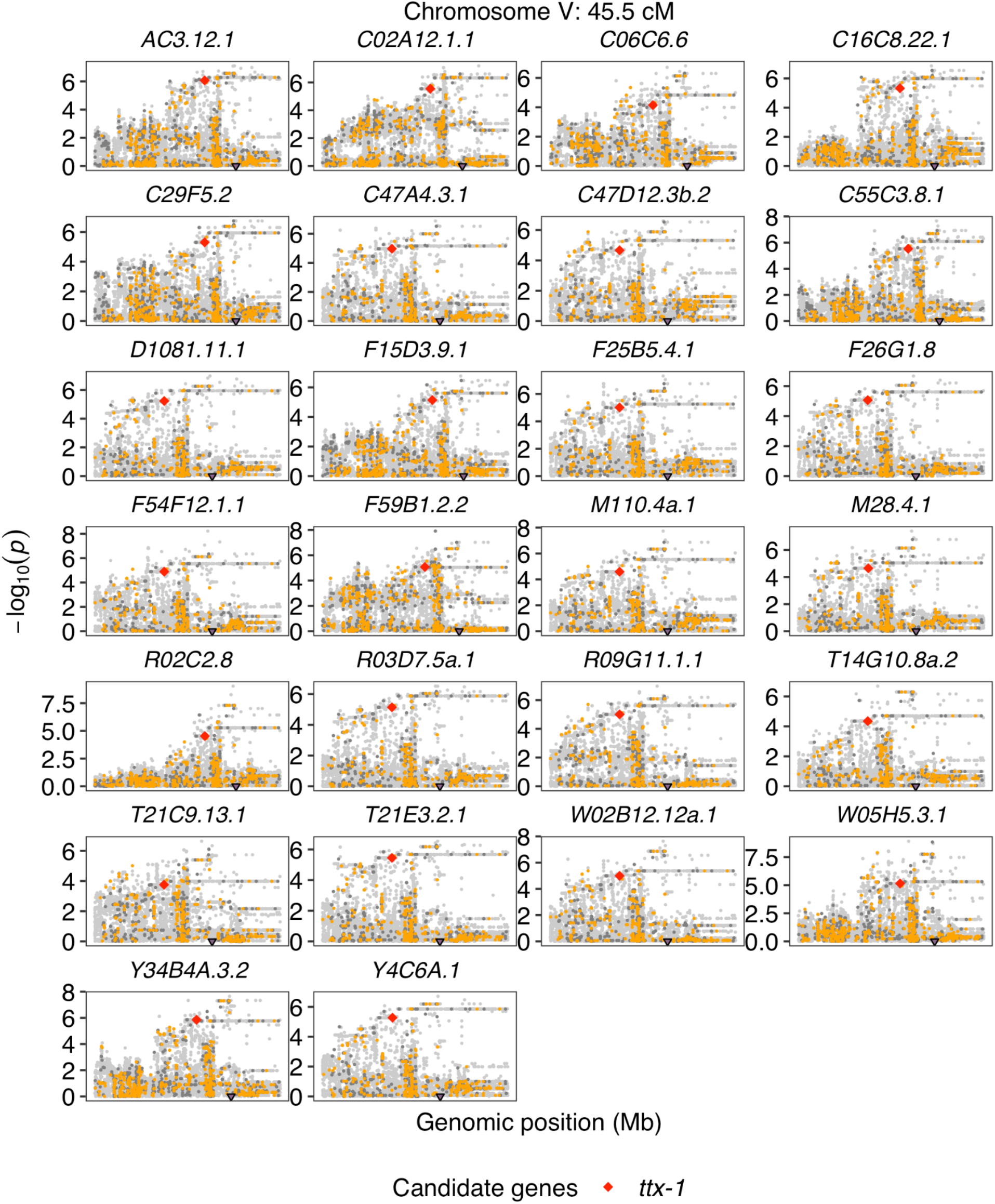
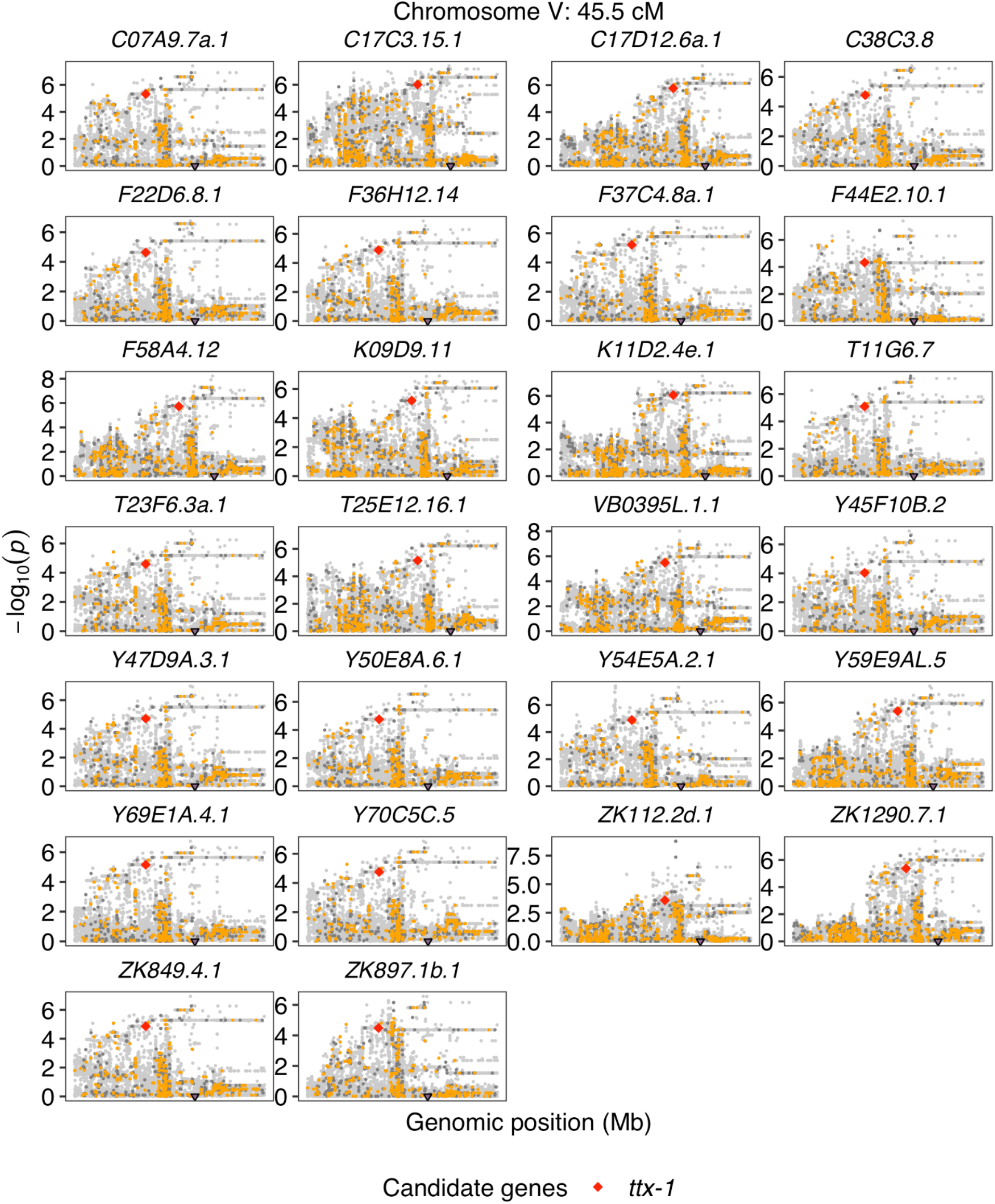
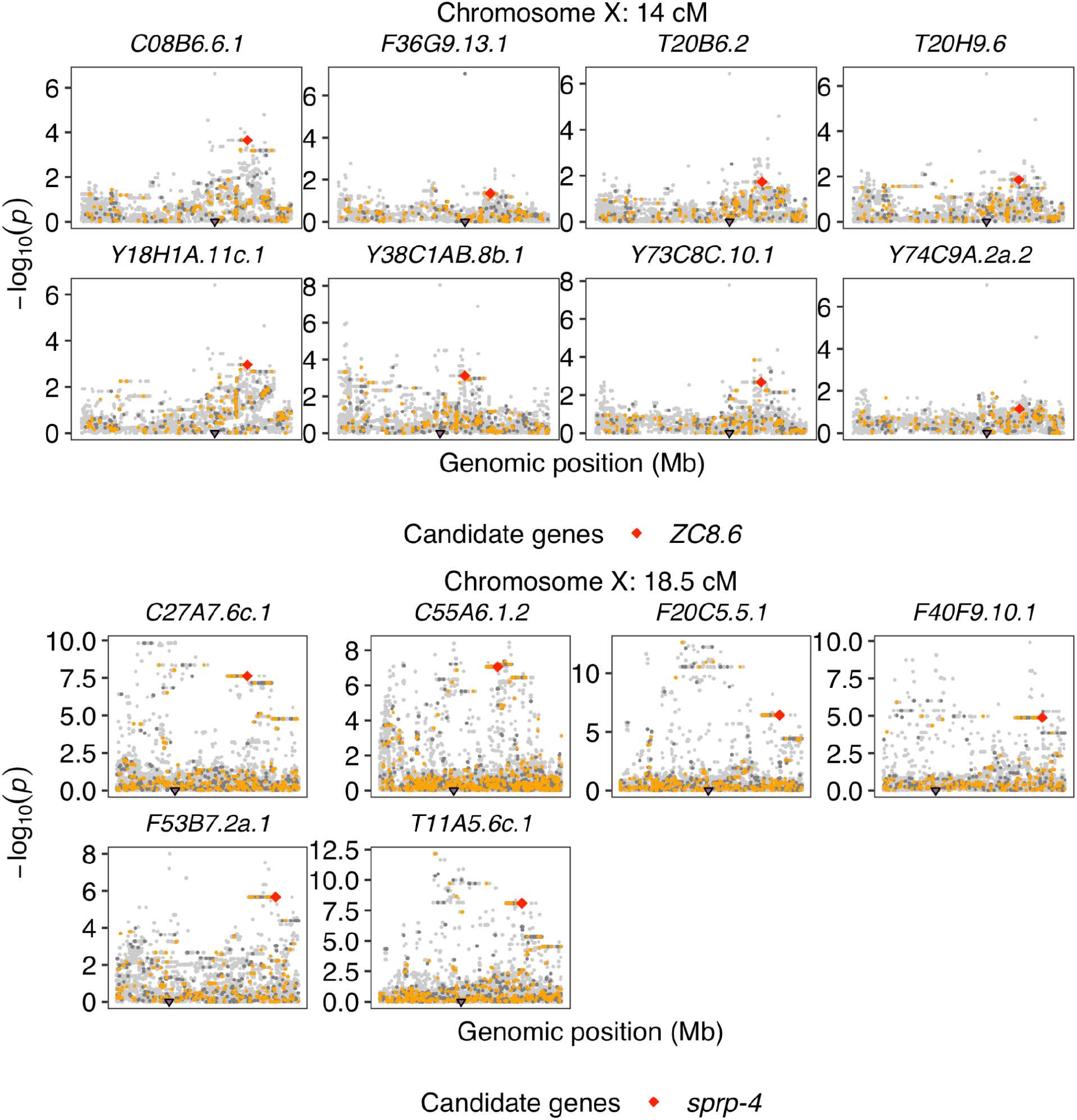
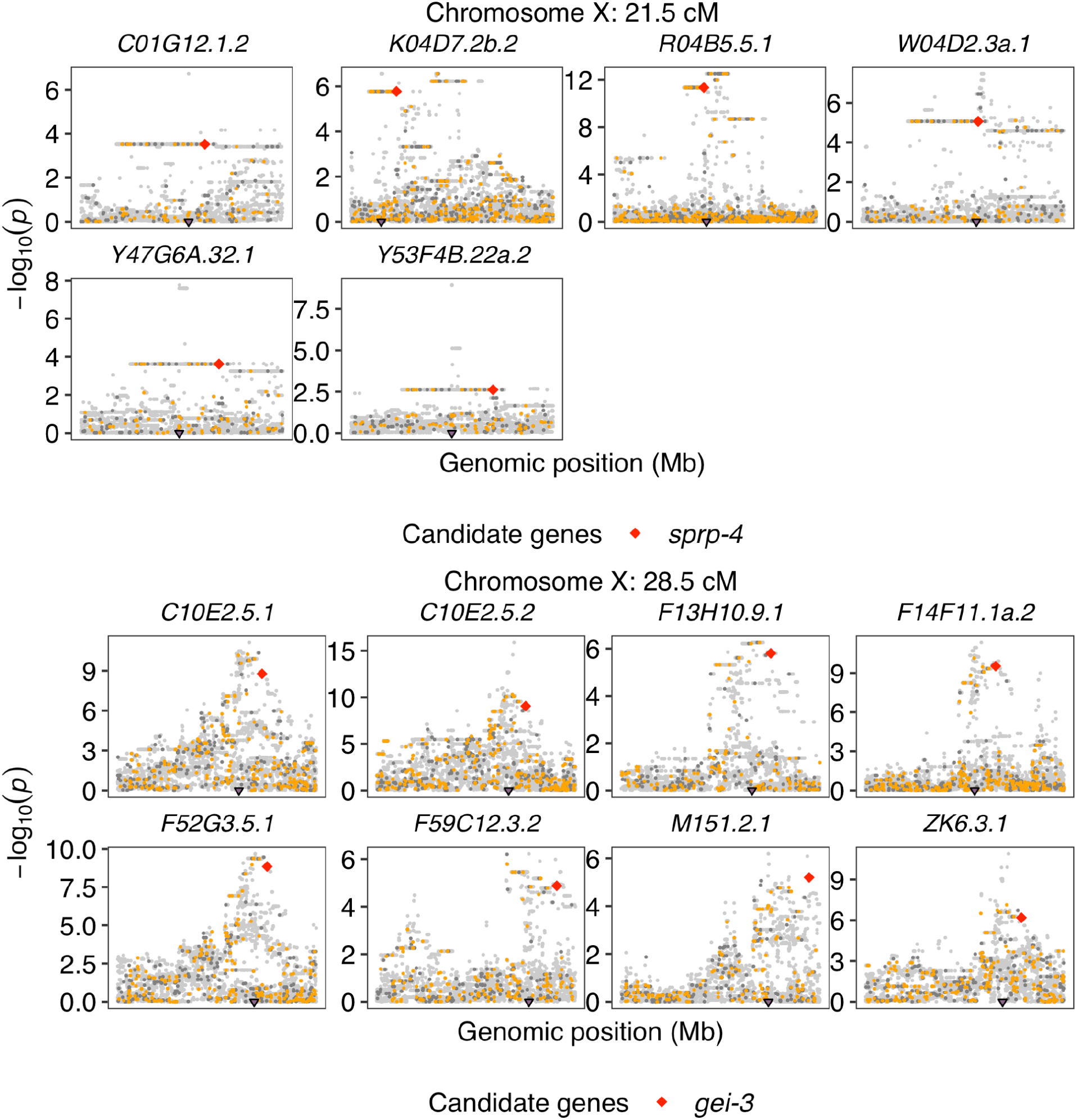
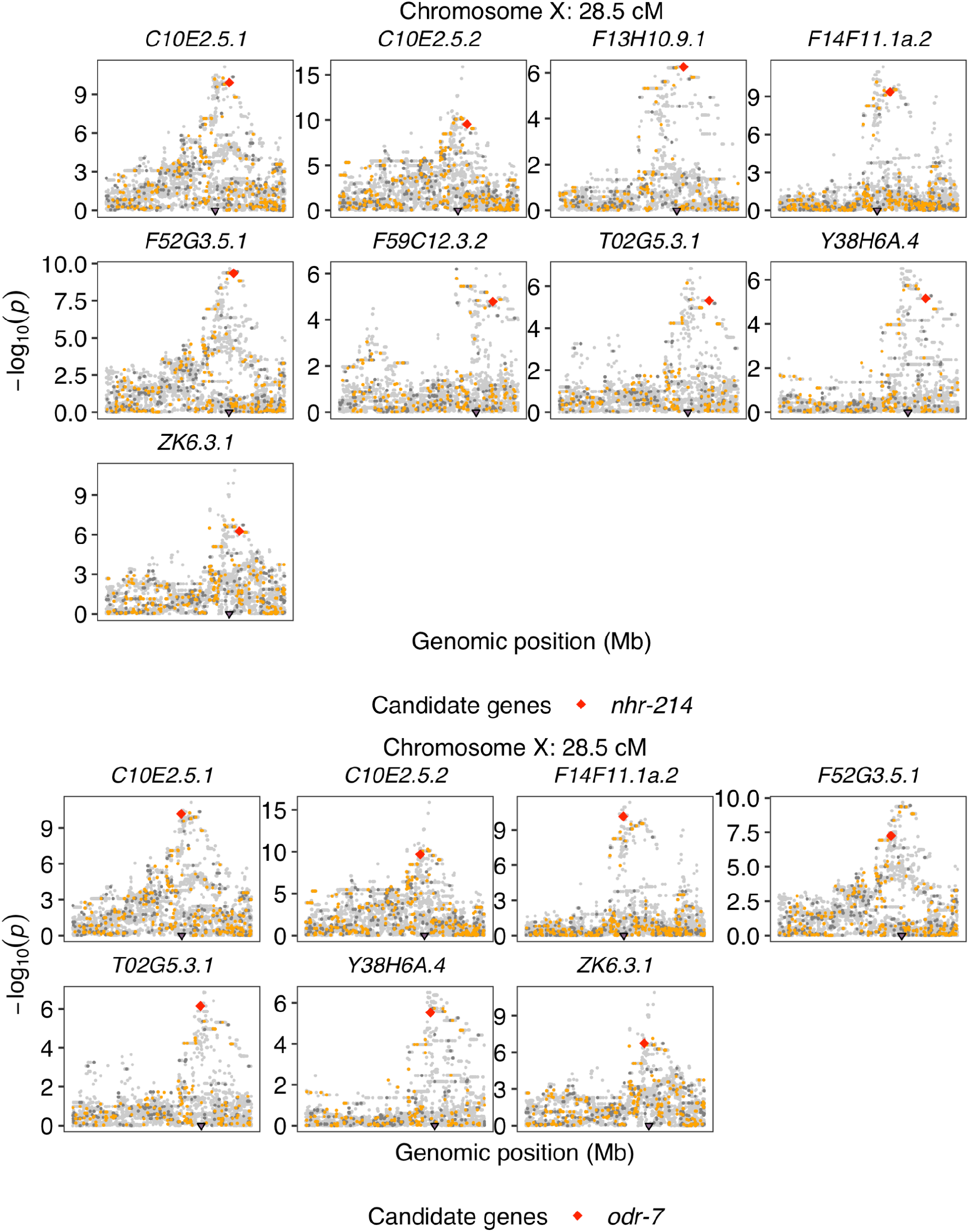
Number of genes encoding chromatin cofactors and transcription factors in each 0.5 cM bin of the *C. elegans* genome. Bins that were identified as distant eQTL hotspots are colored red. Other bins are colored black.

**Supplementary Fig. 7.**
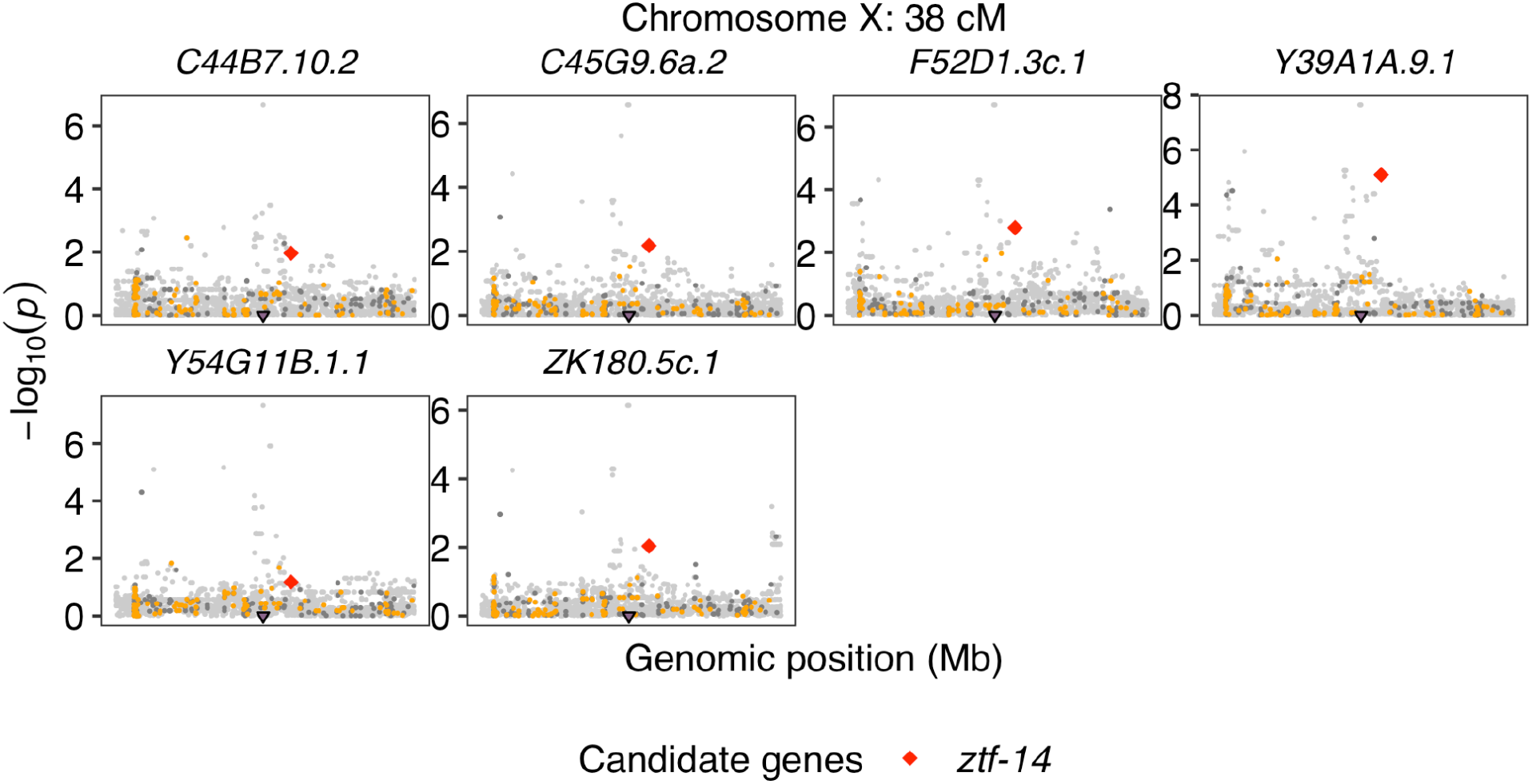
Fine mapping of transcript expression traits with distant eQTL in different hotspots is shown. Genomic position (x-axis) is plotted against the -log10(*p*) values (y-axis) for each variant. Purple triangles on the x-axis represent eQTL positions. Candidate variants with negative BLOSUM scores in genes encoding transcription factors or chromatin cofactors are indicated as red diamonds. Other variants that are with negative BLOSUM scores, with non-negative BLOSUM scores or intergenic are colored orange, dark gray, and light gray, respectively. Transcript names of each trait are indicated above each panel.

**Supplementary Fig. 8.**
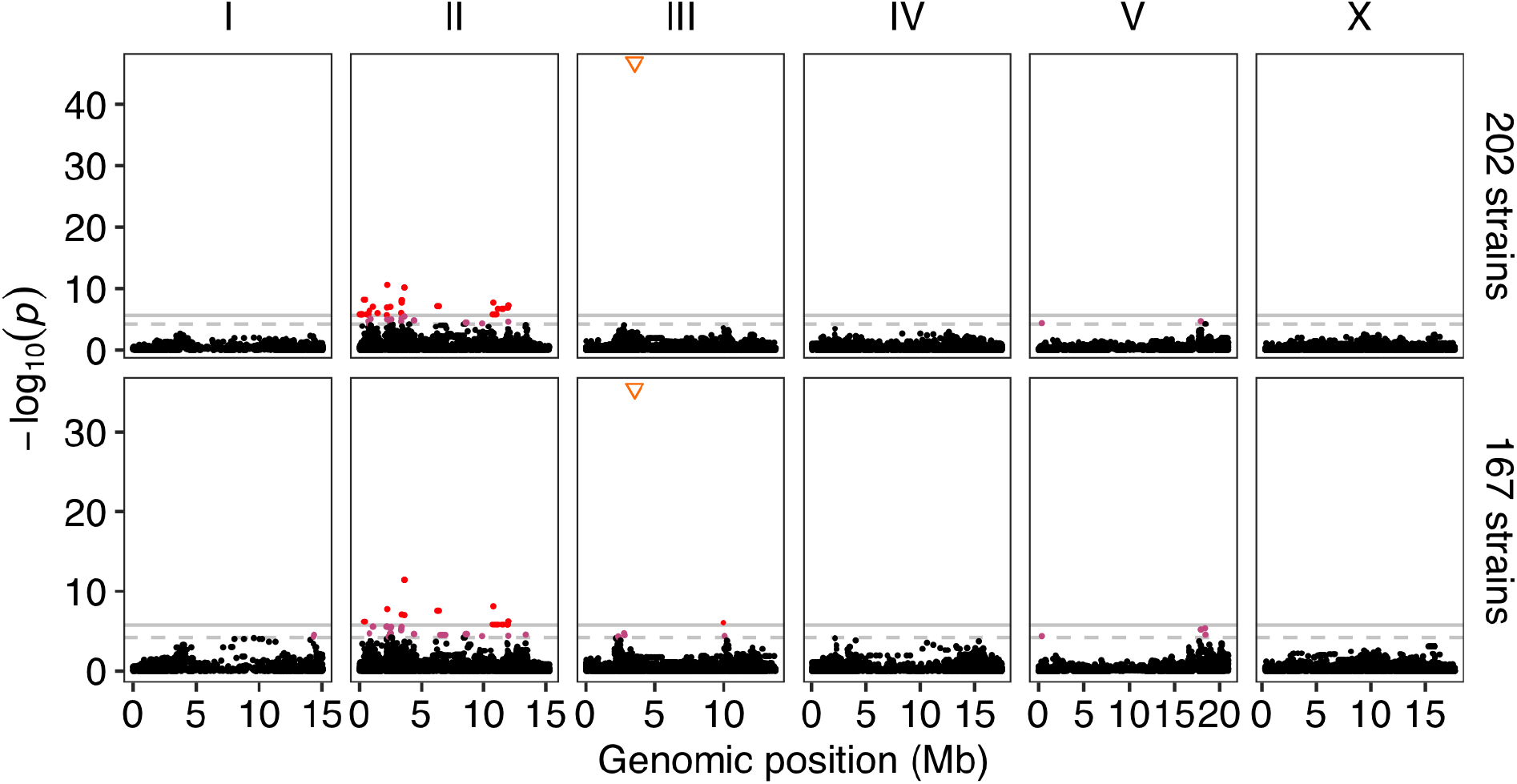
Manhattan plots indicating the GWA mapping result for animal length (q90.TOF) of 202 (top panel) and 167 (bottom panel) *C. elegans* wild strains in response to ABZ^44^ are shown. Each point represents an SNV that is plotted with its genomic position (x-axis) against its -log_10_(*p*) value (y-axis) from the GWA mapping. Real SNVs that pass the genome-wide EIGEN threshold (the dotted gray horizontal line) and the genome-wide Bonferroni threshold (the solid gray horizontal line) are colored pink and red, respectively. The pseudo SNV marker representing high allelic heterogeneity in the gene *ben-1* at position 3,539,640 on chromosome III is indicated as an orange inverted triangle.

**Supplementary Fig. 9.**
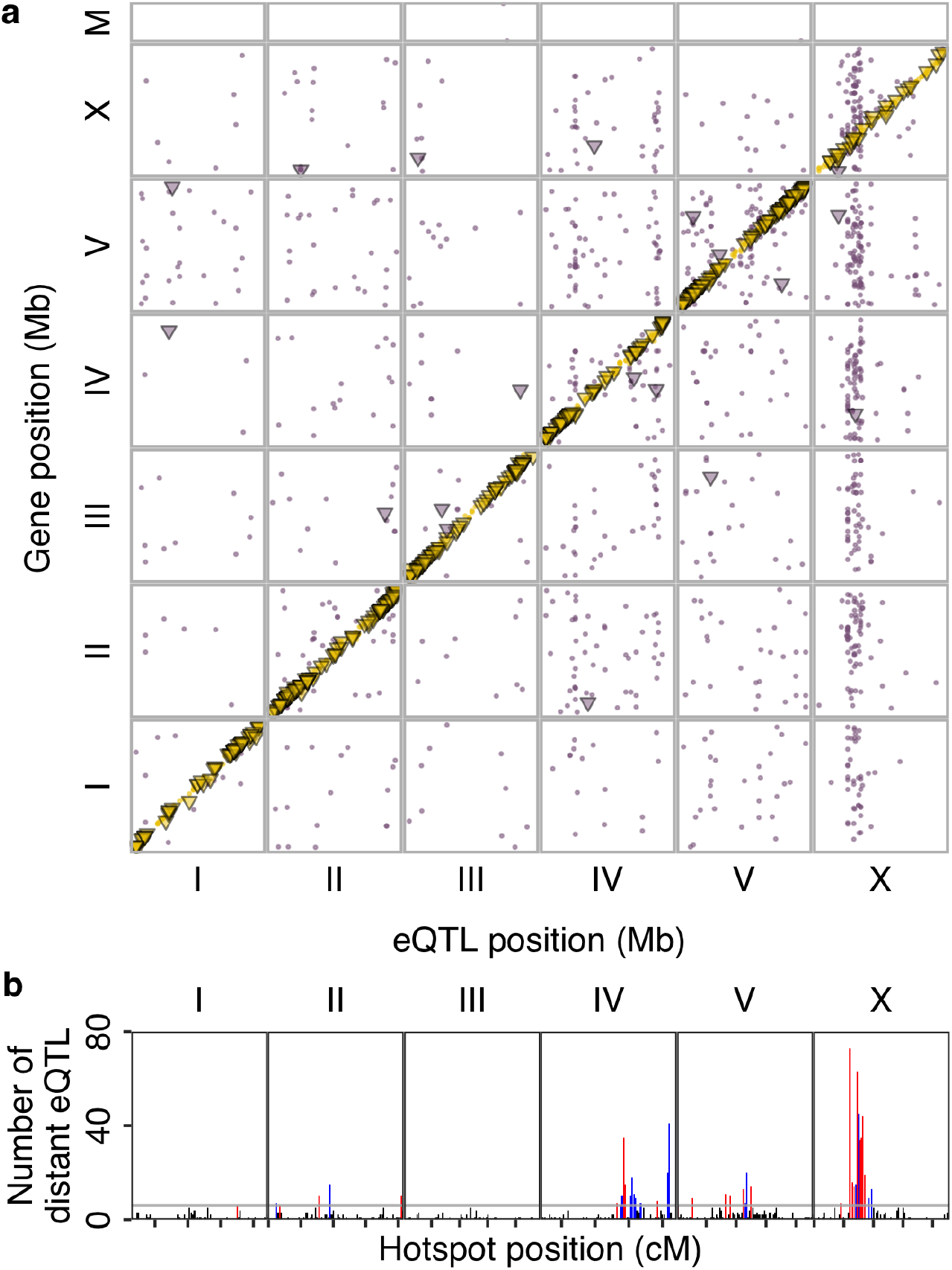
**RIAILs eQTL. a**, The genomic locations of 2,387 eQTL peaks (x-axis) in the RIAILs eQTL studies^3, 9^ are plotted against the genomic locations of the 2,003 genes with expression differences (y-axis). Golden points or triangles on the diagonal of the map represent local eQTL. Purple points or triangles correspond to distant eQTL. Triangles represent eQTL that were also found in our study. **b**, The number of distant eQTL (y-axis) in each 0.5 cM bin across the genome (x-axis) is shown. Tick marks on the x-axis denote every 10 cM. The horizontal gray line indicates the threshold of 6 eQTL. Bins with 6 or more eQTL were identified as hotspots and are colored red or blue. Bins with fewer than 6 eQTL are colored black. Blue bins represent hotspots that were also found in our study.

## Notes

### Competing Interest Statement

The authors have declared no competing interest.

